# Loss of Sox10 prevents tumor initiation and induces luminal-to-basal reprograming in a HER2+ mouse model

**DOI:** 10.1101/2025.08.26.672423

**Authors:** Brennan Garland, Samuel Delisle, John Abou-Hamad, Christiano de Souza, Riana Zuccarini, David P. Cook, Rebecca Auer, Luc A. Sabourin

## Abstract

The SRY-HMG-Box transcription factor SOX10 plays a critical role in neural crest development, but its function in epithelial tumorigenesis remains unclear. Here, we identify SOX10 as a key regulator of tumor-initiating activity in Neu-driven mammary cancers. Genetic ablation of Sox10 in the luminal compartment of MMTV-Neu (NIC) mice resulted in delayed but normal mammary gland development. Sox10 deletion resulted in a reduction in mammary progenitors and tumor initiating activity with a complete loss of tumor initiation in Sox10-deficient luminal cells. CRISPR/Cas9-mediated Sox10 inactivation in Neu-transformed tumor cells led to diminished self-renewal in mammosphere assays, markedly impaired growth in orthotopic transplant models and profoundly reduced lung colonization following tail vein injection, suggesting a depletion of cancer stem cell activity. Transcriptomic profiling revealed that Sox10-deficiency in Neu+ tumor cells induces a luminal-to-basal/stem-like shift and the downregulation of several genes associated with genetic networks regulating stemness. Collectively, these findings demonstrate that Sox10 is required for a permissive luminal cell state for Neu-driven tumor initiation and that it is critical for cancer stem cell activity and the establishment of metastases.

**Significance:** The Sox10 transcription factor is critical for mammary stem cell function and plasticity. Animal studies suggest that different subtypes of breast cancers arise from luminal epithelial cells and hypothesized the implication of luminal stem cells in breast cancer initiation. We show that Sox10 deletion in the luminal compartment of adult female mice abrogates tumor initiation. Sox10 inactivation impairs tumor cell growth, dissemination and reprograms luminal Neu+ tumor cells to a basal phenotype. Our data supports a model whereby luminal progenitors are required for tumor initiation, self-renewal and growth at distant sites. The findings suggest that therapies aimed at interfering with Sox10 activity or expression might be beneficial in the treatment of breast cancers.

## Introduction

The HER2/ErbB2/Neu receptor tyrosine-kinase (RTK) is overexpressed in approximately 25-30% of human breast cancers (1, 2). Currently, the most effective form of treatment for HER2+ breast cancer remains chemotherapy in combination with trastuzumab (Herceptin). However, acquired or *de novo* resistance is observed in up to 70% of HER2+ metastatic breast cancers (3, 4). Murine models of HER2-positive breast cancer have confirmed the oncogenic role of HER2 in breast tumor initiation and progression (5, 6), yet identification of the cell population susceptible to HER2-mediated transformation remains largely undefined. This highlights the need to investigate regulators of lineage plasticity as a potential prerequisite for malignant reprogramming.

Sox10 (Sry-HMG Box 10) is a critical transcriptional regulator of cell fate determination and several developmental processes during embryogenesis (7, 8). Recently, it has been found that Sox10 expression is conserved in mammary epithelial cell populations, specifically in those with the highest degree of stem/progenitor cell activity (9). Consistent with this, conditional deletion of Sox10 in mice has been shown to disrupt normal mammary gland development, further highlighting its functional role in the maintenance of the mammary epithelial lineage (10).

Currently, the role of Sox10 in malignant progression remains elusive and controversial with different observations in various types of cancers (11, 12). In the context of breast cancer, we and others have shown that Sox10 is a marker for select basal-like carcinomas such as triple-negative breast cancers (13–17). Recenly, Sox10 has been shown to be highly expressed in luminal progenitors of the human mammary glands and is over-represented in TNBC brain metastases (18).

Using murine models of luminal-, and basal-like mammary tumorigenesis, Sox10-expressing tumours exhibited a de-differentiated state and acquired a more mesenchymal-like phenotype (19). Mechanistically, Sox10 was found to be directly bound to promoter regions of genes regulating epithelial-mesenchymal transition (EMT), stem/progenitor cell activity, and neural crest identity (19). Functional studies using C3-TAg mice, a model of basal-like breast cancer, revealed that Sox10 haploinsufficiency significantly prolongs survival, directly implicating it in disease progression (19). Supporting this, using a murine model of HER2+ breast cancer, we have shown that increased numbers of Sox10+ cells correlates with a more rapid tumor initiation (20).

Our lab has recently shown that TCGA datasets revealed that approximately 15% of HER2+ breast cancers are Sox10^hi^ (21). Consistent with this, others have demonstrated substantial variability in proportions of the Sox10+ populations (2% - 79%) among HER2+ tumour samples (20, 22–25). Interestingly, Sox10^hi^ cohorts were associated with high EMT-like signatures, worse grade, and a stem-like phenotype, suggesting that Sox10+/HER2+ tumors represent a more aggressive de-differentiated subtype (20, 22, 25).

Despite these observations, whether the Sox10+ luminal population is critical for HER2-driven tumorigenesis or represents a transitional state of cellular plasticity remains unresolved. Here, we investigated the functional and genetic consequences of conditional Sox10 deletion in MMTV-NIC mice (26). We find that luminal deletion of Sox10 delays mammary gland development but completely abrogate hyperplasia development and tumor initiation in MMTV-NIC mice. CRISPR-mediated deletion of Sox10 in established Neu+ cell lines impairs tumor growth in orthotopic assays and prevents tumor expansion in lung colonization studies. Tumor cells devoid of Sox10 show a luminal to basal/stem-like shift in gene signature. These results suggest that Sox10+ luminal progenitors are required for Neu-induced tumorigenesis and that the loss of Sox10 can reprogram cancer cells to a basal-like state.

## Results

### Sox10 inactivation delays mammary gland development

Previous studies have shown that Sox10 is required for full development of the mammary gland (10). However, the MMTV-Cre strains used have been shown to display intrinsic developmental defects due to transgene positional effects and leaky expression in the myoepithelial/basal layer (27, 28). As the MMTV-NIC and MMTV-Cre lines used here have been shown to be exclusively expressed in the luminal compartment (26, 29–31), we first assessed mammary gland development in this background.

Sox10*^fl/fl^* mice (32) were crossed with MMTV-Cre mice (29) and mammary gland development was assessed at 4 and 11 weeks of age. As previously reported (29), in a MMTV-Cre:ROSA26 background, LacZ staining marked Cre expression in rudimentary mammary glands as early as P14 (Suppl. Fig.1A). At 4-weeks of age, MMTV*-*Cre:Sox10*^fl/fl^*females showed a slight delay in mammary tree invasion, end bud numbers and ductal branch point morphogenesis (Fig. 1A-C). However, by about 11-weeks of age, no obvious defects were observed in post-pubertal virgin females and mammary glands were indistinguishable from that of the wildtype group (Fig.1A-C). Furthermore, no defects in epithelial organization were observed (Fig.1D) and similar proportions of luminal (CK8) and basal cells (SMA) were present (Suppl. Fig. 1B,C). While basal cells remained Sox10+, immunofluorescence showed lower expression of luminal Sox10 and approximately a 60% reduction in Sox10+ luminal cells in 11-week-old in MMTV*-*Cre:Sox10*^fl/fl^* females (Fig. 1E, F). Inefficient but progressive deletion of this Sox10 allele throughout the lifespan of the animals has also been observed previously (32). Supporting this, in 16 week-old MMTV-NIC mice, other than the basal layer, virtually all luminal cells were devoid of Sox10 (Suppl. Fig.1D,E). In contrast to a previous report (10), it is likely that delayed inactivation of Sox10 in luminal progenitors in our model resulted in the development of mature mammary glands.

**Figure 1.**
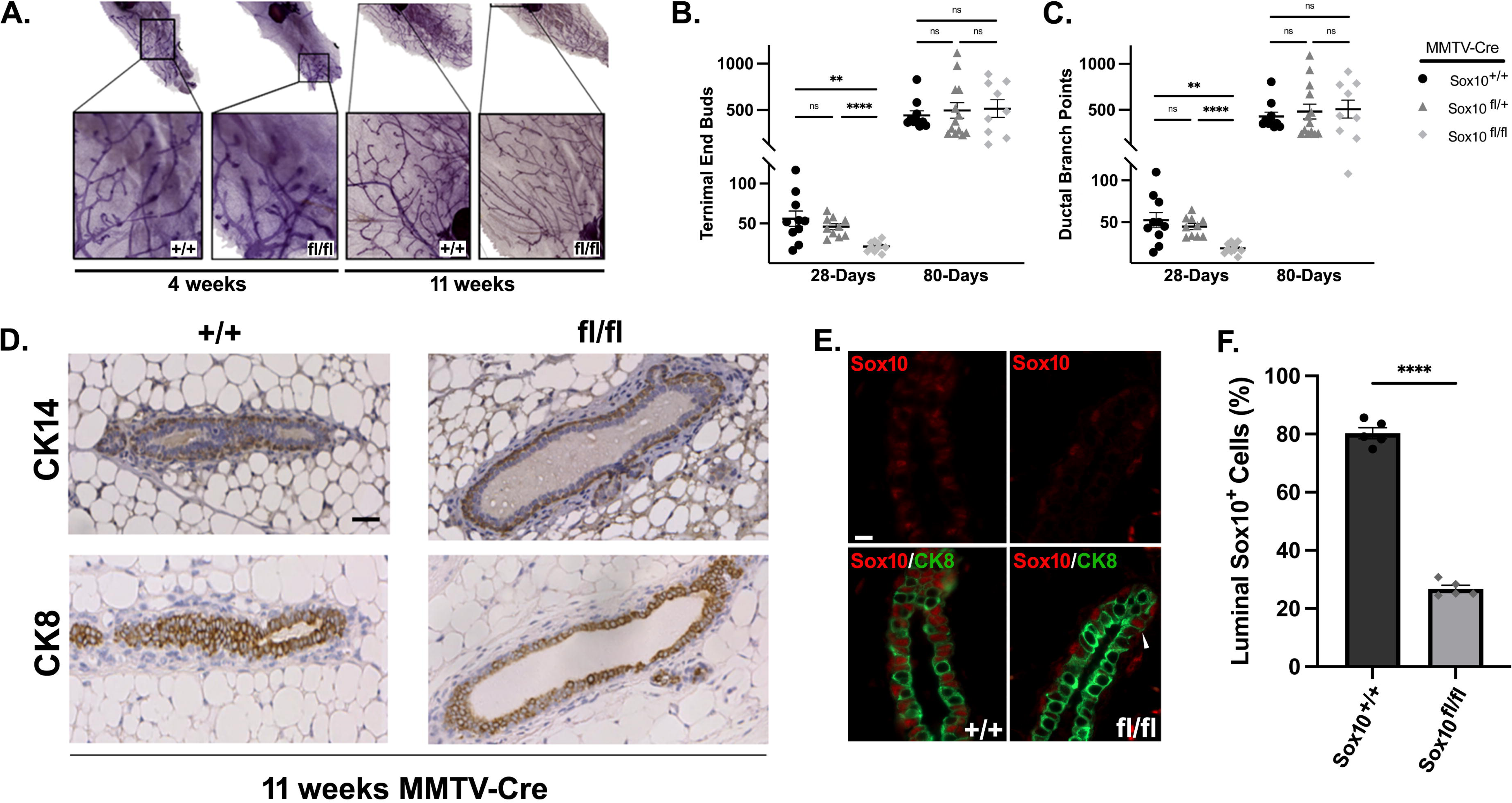
Luminal *Sox10*-deletion delays mammary gland development. (A) Representative images of hematoxylin stained MMTV-Cre:Sox10^+/+^ and MMTV-Cre:Sox10^fl/fl^ mammary gland whole-mounts at 4-, and 11-weeks of age. Quantification of epithelial end buds (B) and ductal branch points (C) in stained MMTV-Cre:Sox10^+/+^; Sox10^+/fl^ and Sox10^fl/fl^. Data shown as mean + SEM. n > 10 mice/group. (D) Immunohistochemistry on MMTV-Cre:Sox10^+/+^ and MMTV-Cre:Sox10^fl/fl^ mammary glands at 11 weeks of age showing similar proportions of luminal (CK8) and basal (CK14) cells. See also Suppl.Fig.1. (E) MMTV-Cre:Sox10^+/+^ and MMTV-Cre:Sox10^fl/fl^ glands were stained for Sox10 expression and CK8 at 11 weeks of age and the efficiency of luminal Sox10 deletion was quantitated in panel (F). Arrowhead shows a residual Sox10+ luminal cell. Scale bar 20μ. **,****p < 0.05, and 0.0005, respectively.

### Sox10 deletion in MMTV-NIC mice inhibits tumor initiation

To assess the role of Sox10 in Neu-mediated mammary tumourigenesis, Sox10*^fl/fl^* mice were crossed with transgenic MMTV-Neu-IRES-Cre (MMTV-NIC) mice (26), driving luminal-specific expression of a bicistronic transcript containing both a constitutively active Neu/ErbB2 (NeuNDL2-5) and Cre recombinase (29). Kaplan-Meier analyses showed a significant increase in disease-free survival (average 76-days) for MMTV-NIC:Sox10*^fl/+^* heterozygote mice when compared to wild-type MMTV-NIC:Sox10^+*/+*^ (Fig. 2A). Surprisingly, MMTV-NIC:Sox10*^fl/fl^* mice remained disease free for the duration of the study, suggesting a complete absence of tumor initiation. Western blot and densitometry analysis of tumour lysates from wildtype and MMTV-NIC:Sox10*^fl/+^*mice revealed similar levels of Sox10 expression along with the luminal marker CK8 (Fig. 2B; see Suppl. Fig.8A). Although some variability was observed, no significant differences were found in the levels of pAKT or pERK1/2, suggesting that MAPK and PI3K signaling downstream of Neu is unaffected *in vivo*.

**Figure 2.**
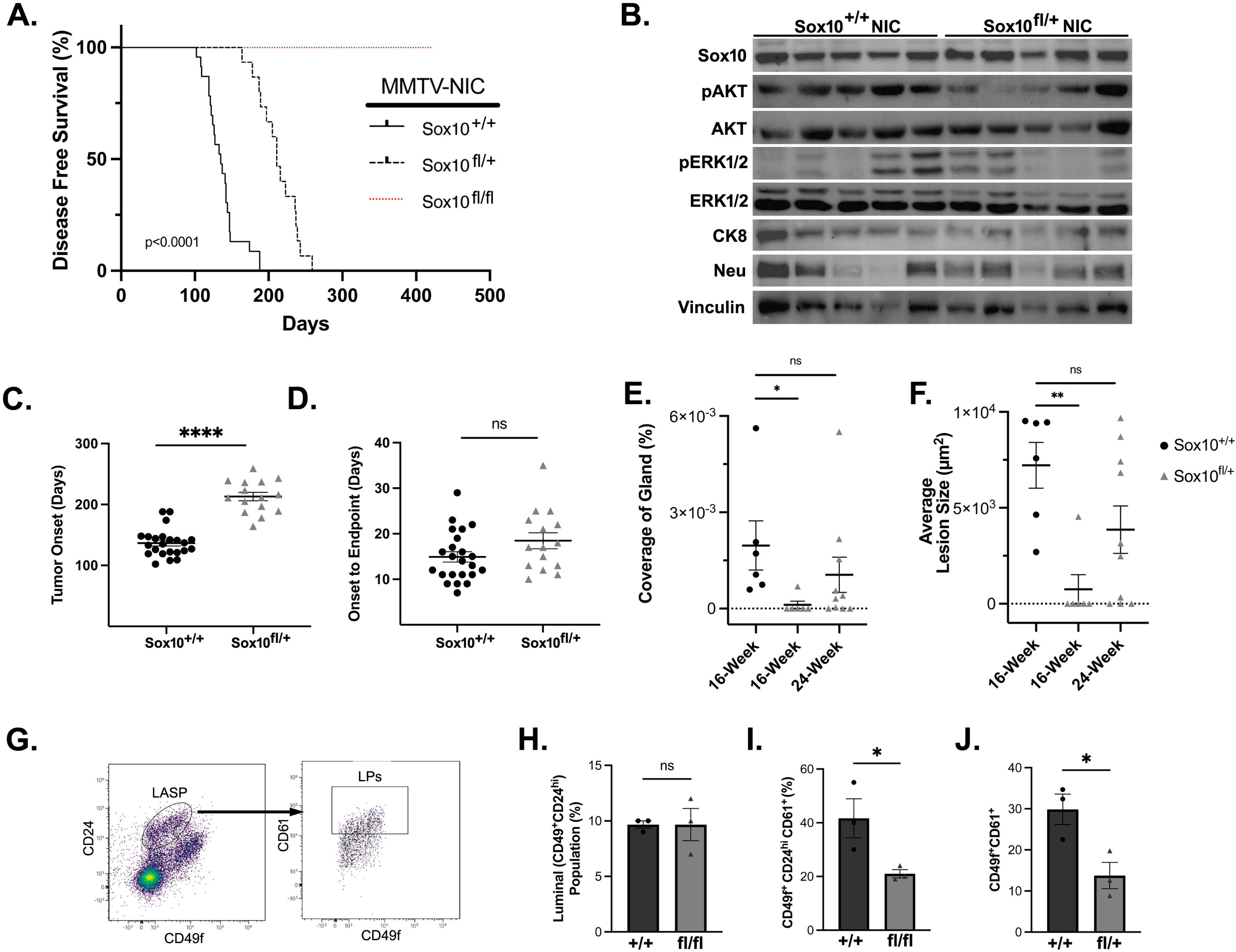
Luminal *Sox10*-deletion impairs Neu-induced tumor initiation. **(A)** Kaplan-Meier disease-free survival curves in MMTV-NIC:Sox10^+/+,fl/+,fl/fl^ mice. Significance was assessed by log-rank test. n > 8 mice/group. (**B)** Endpoint tumour lysates were assessed for the expression of Sox10, Neu, pAKT (S473), pERK1/2 (Y204) and CK8 using Western blot analysis. Time for tumour onset (0.5 cm^3^) (**C**) and progression to endpoint (1.5 cm^3^) (**D**) were measured for all MMTV-NIC genotypes. n> 15 mice/group. Quantification of total mammary gland coverage (E) and average size of hyperplastic lesions (F) in 16-, and 24-week-old MMTV-NIC mice from all genotypes. n> 6 mice/group. (G) Flow sorting strategy for measurements of the LASP compartment. Lin-cells were first sorted into the luminal population (CD24/CD49f; circled) and further quantitated using CD61/CD49f. (H) Quantitation of the luminal population (CD49f^+^/CD24^hi^) in 8-week old MMTV-Cre females. (n=3) (I) Quantitation of the CD24^hi^/CD49f^+^/CD61^+^ populations in 8-week old MMTV-Cre females (n=3) (J) Quantitation of the CD49f^+^/CD61^+^ populations in 16-week old MMTV-NIC:Sox10^fl/+^ females (n=3). Data shown as mean ± SEM. *,**,****p < 0.05,0.01, and 0.0005, respectively.

Palpable tumours were readily detectable in MMTV-NIC:Sox10^+/+^ about 80 days sooner than in MMTV-NIC:Sox10*^fl/+^* mice (Fig.2C). However, time from onset to endpoint was unchanged in either group, suggesting that the growth of established tumors was unaffected (Fig. 2D). Supporting our onset to endpoint data, no quantitative differences were observed for the proliferative and apoptotic markers, Ki-67 and cleaved caspase-3, respectively (Suppl. Fig.2A-D).Although Sox10 expression was similar in endpoint tumors (Fig.2B), one possibility is that Sox10 haploinsufficiency in non-transformed luminal cells significantly delays tumor initiation. To further investigate the observed delay in tumour onset, mammary glands were collected at 16-weeks of age, prior to tumor detection, and assessed for hyperplastic lesions. Sox10 staining in 16 week-old MMTV-NIC:Sox10*^fl/fl^*revealed a marked reduction in Sox10+ luminal cells with numerous SMA+/Sox10+ basal cells (see Suppl. Fig.1,E), suggesting efficient luminal Sox10 excision. Hematoxylin staining of 16-week-old mammary gland sections revealed the presence of numerous hyperplastic lesions in wildtype MMTV-NIC:Sox10^+*/+*^ mice (Fig.2E,F; Suppl. Fig.2G). However, no hyperplasia were observed in age matched MMTV-NIC: Sox10*^fl/+^*mice (Fig.2E,F; Suppl. Fig.2H). On average, an additional 8 weeks was required for heterozygote mice to display hyperplastic lesions detectable by haematoxylin stain (Fig.2E,F; Suppl. Fig.2I). Supporting an absence of tumor formation, no hyperplasia was ever detected in MMTV-NIC: Sox10*^fl/fl^* mice, up to 420 days of age (Suppl. Fig.2J). Together, these data suggest that tumor initiation is abrogated in Sox10-deficient luminal cells.

The term LASP (luminal adaptive secretory precursor) has been proposed to describe the most disparate group of mammary luminal cells encompassing luminal/alveolar progenitors/precursors, luminal secretory and hormone receptor-negative (33). We will hereafter use LASPs to refer to luminal cells with potential stem/progenitor activity. Genetic studies revealed that Sox10 is expressed in a progenitor subset of LASPs (9, 18, 34–37). Because of the critical role of Sox10 in murine mammary stem/progenitor cell function (9, 19), we assessed the LASP compartment using flow cytometry (38–43). Flow sorting of the Lin^-^/CD24^hi^/CD49f^+^ in 8 week-old MMTV-Cre virgin females showed no difference in the total luminal populations between wild type and Sox10^fl/fl^ mice (Fig.2G,H and Suppl. Fig8.C). However, further sorting of the luminal compartment into CD24^hi^/CD49f^+^/CD61^+^, enriched for progenitor activity (41–43), shows a 2-fold reduction in the LASP population in Sox10-deficient mammary glands (Fig.2I and Suppl. Fig.8D). In transgenic models of spontaneous mammary tumors, the Lin^-^/CD49f^+^/CD61^+^ population was found to be enriched for tumor initiating activity (44, 45). Supporting this, a two-fold reduction in the Lin-/CD49f^+^/CD61^+^ population was observed in 16 week-old MMTV-NIC:Sox10*^fl/+^* heterozygote mice, suggesting a depletion of tumor initiating activity upon the loss of a single copy of Sox10 (Fig.2J and Suppl. Fig.8E).

The Sox9 transcription factor has been shown to be critical for luminal lineage commitment and tumor progression in a TNBC mouse model (36, 46, 47). Therefore, we assessed Sox9 expression in 16 week-old mammary glands. For all MMTV-NIC:Sox10 genotypes, strong Sox9 expression was observed in both the basal and luminal layers (Suppl. Fig.2K), suggesting that the loss of Sox10 does not affect Sox9 expression. Importantly, despite Sox9 expression, tumor initiation was impaired suggesting that Sox9 cannot compensate for Sox10 in this process. Together, these data suggest a critical role for Sox10 in the maintenance of LASPs and in Neu-induced mammary tumourigenesis.

### Sox10-deficient Neu^+^ cell lines have reduced tumorigenic potential in vivo

Immunohistochemical analyses of endpoint tumors revealed that all tumors from wildtype and MMTV-NIC:Sox10*^fl/+^* heterozygotes expressed Sox10 and histologically resembled solid/nodular adenocarcinomas (6, 26) (Suppl. Fig.2E,F). However, as previously reported for human HER2L tumors (20, 22, 23, 25), the number of Sox10+ nuclei was found to vary greatly between tumors (∼15-80%; see Suppl. Fig.2F), potentially reflecting the heterogeneity and the variable CSC content of those tumors. Similarly, western blot analysis of MMTV-NIC- and MMTV-NDL-derived tumor cell lines showed markedly different levels of Sox10 expression (Suppl. Fig.3A).

To further characterize the observed *in vivo* phenotype, we derived new low passage Sox10-expressing tumor cell lines from endpoint tumors. Two independent tumor cell lines expressing Sox10 (BG1 and BG2; Suppl. Fig.3A) were successfully established from MMTV-NeuNDL tumour-bearing mice (5, 48). Immunostaining showed on average approximately 75 and 90% *Sox10*+ nuclei for BG2 (Fig.3A) and BG1 cells (Suppl. Fig.3D), respectively. To inactivate Sox10, two guide RNAs (sgRNA/RFP) targeting amino acids 168-175 (sgRNA-1) or 67-74 (sgRNA-2) were transduced into cell lines expressing Cas9 and RFP+ cells were flow sorted. Sox10 knockout was then assessed by immunofluorescence, western blotting and RT-qPCR. In both lines, transduction with sgRNA-2 resulted in markedly reduced levels of Sox10 protein and less than 5% of *Sox10*+ nuclei (see Fig. 3A, B and Suppl. Fig.3C). Therefore, the sgRNA-2 pools were selected for further studies. Both Neu+/Sox10^-^ BG1 and BG2 cell lines exhibited growth rates similar to that of the non-targeting control (NTC) in culture (Suppl. Fig.3E,F), suggesting that Sox10 is not required for tumor cell proliferation or cell viability *in vitro*. Furthermore, the Sox10-deficient cell lines retained Neu expression, suggesting that the transgene is not affected by the loss of Sox10. Although, no differences were observed in the levels of pERK1/2, higher levels of pAKT were detected in the two BG1 Sox10-null clones (Suppl. Fig.3C and 8B).

**Figure 3.**
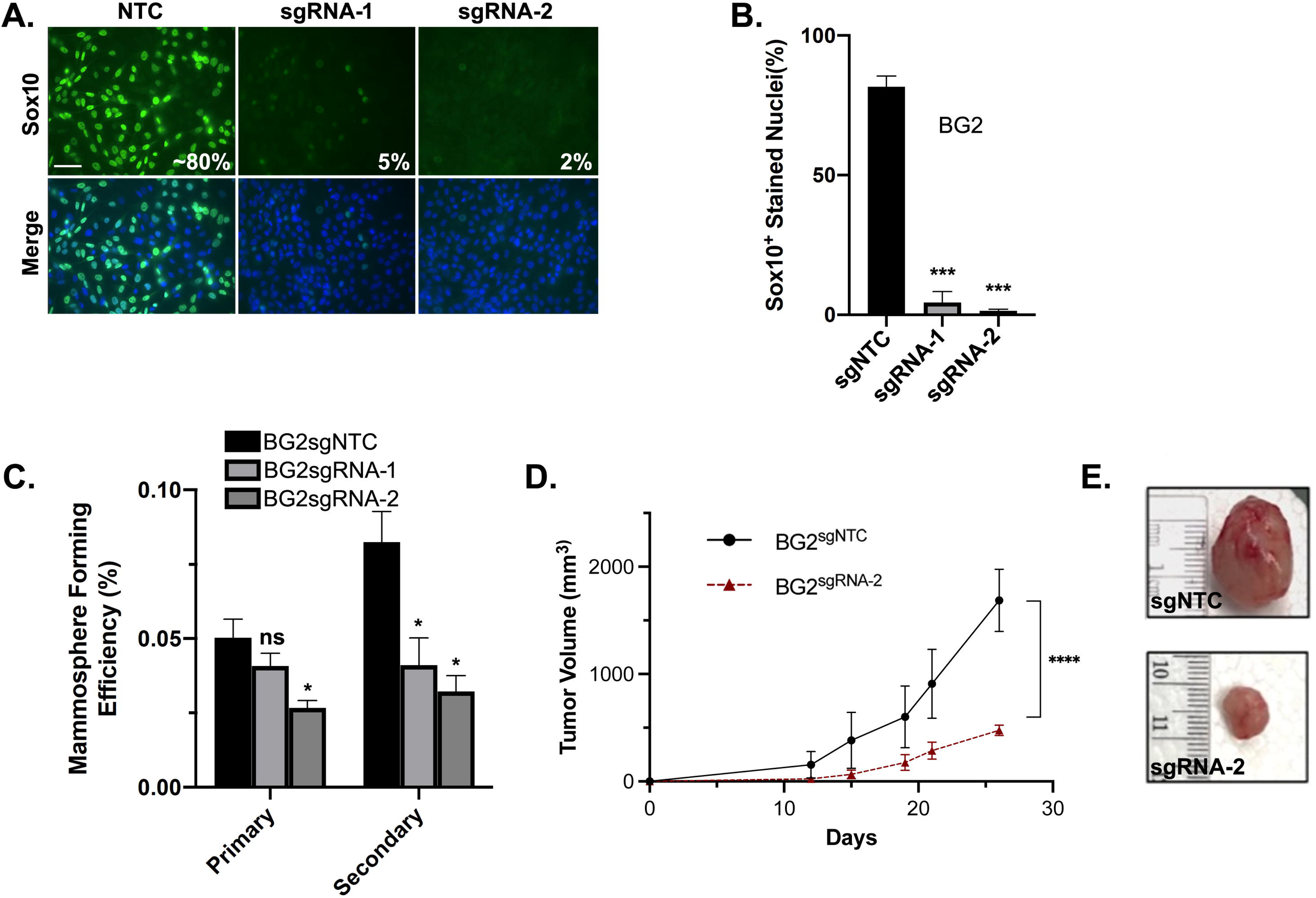
*Sox10*-deficient NeuNDL cell lines have reduced stemness and tumorigenic potential. (A) Immunofluorescence and DAPI staining for Sox10 (green) and (B) proportions of Sox10+ nuclei following CRISPR-Cas9 mediated *Sox10* ablation in NeuNDL BG2 cells using two independent guide RNAs. Scale bar 50 μ. (**C**) Primary and secondary mammosphere efficiencies were measured in BG2-sgRNA-1, sgRNA-2 and NTC pools after 7-days in ultra-low attachment plates and imaging. (D) 10^6^ BG2-sgRNA-2 or NTC cells were injected orthotopically into NCG mice and tumour volumes were recorded bi-weekly until an endpoint of 1.7cm^3^ was reached by the first tumour in the cohort. n = 4 mice/group. (**E**) Examples of resected tumors from cells injected in (D). Data shown as mean + SEM. *,***,****p < 0.05, 0,01 and 0.0005, respectively.

Deletion of Sox10 did not have any effect on cell migration as tested in transwell chambers (Suppl. Fig.3G,H). Supporting previous data in mammary organoids (9), Sox10-deficient cells showed a marked reduction in secondary mammosphere formation, suggesting impaired stem/progenitor activity (Fig. 3D and Suppl. Fig.3I). For all assays, both Sox10-deficient lines showed similar results. Although no effect on proliferation was observed *in vitro*, orthotopic injection of pooled BG2-Sox10^+^ and Sox10^−^ tumor cells into the mammary fat pad of immunodeficient mice revealed a marked delay in tumor growth in the Sox10-deficient group (Fig. 3D, E). Interestingly, Sox10 immunohistochemistry on tumors at endpoint showed that the proportion of Sox10+ cells was similar between Sox10+ and Sox10^-^ cell lines (Suppl. Fig.3J). As the pool of BG2-Sox10^−^ injected cells did not show a complete loss of Sox10 (see Fig.3A), it is likely that the tumors arising upon injection of Sox10^-^ cells are due to residual Sox10-expressing subpopulations. As Sox10 deletion does not affect cell proliferation *in vitro*, these data suggest that Sox10 loss impairs tumorigenic potential *in vivo*.

Western blot analyses show a wide range of Sox10 expression in NIC- and Neu-NDL-derived tumor lines with some showing undetectable levels (see Suppl. Fig.3A), suggesting that Sox10 expression may be lost upon prolonged passaging in culture. Supporting this, we have observed *Sox10* loss over time in BG2 cells (see Suppl. Fig.3B). However, BG1 cells displayed stable Sox10 expression over multiple passages and were selected for additional *in vivo* studies. To avoid the potential expansion of residual Sox10**^+^**cells upon transplantation, the BG1sgRNA-2 pools were cloned and expanded. Two Sox10-deficient clones were randomly selected for further studies (see Suppl. Fig.3C). For both clones, sequencing analyses showed deletions or mutation of amino acids 76 and 77, resulting in premature stop codons in exon 4.

To compare the tumorigenic potential and cancer stem cell (CSC) activity, ten-fold dilutions of BG1Sox10**^+^** and Sox10**^-^**cells were injected orthotopically in NSG mice. Tumor volume measurements over time showed no measurable growth in all animals injected with BG1 Sox10-deficient tumor cells (Fig.4A). Supporting this, a marked reduction in tumor weight was observed in all animals bearing Sox10**^-^**cells (Suppl. Fig.4A,B). Other than rare small pockets of Sox10+ cells, immunohistochemical analysis of residual tissue confirmed the absence of *Sox10* expression (Fig. 4B). Together, these data suggest that the CSC activity is markedly impaired in Sox10-deficient cells.

**Figure 4.**
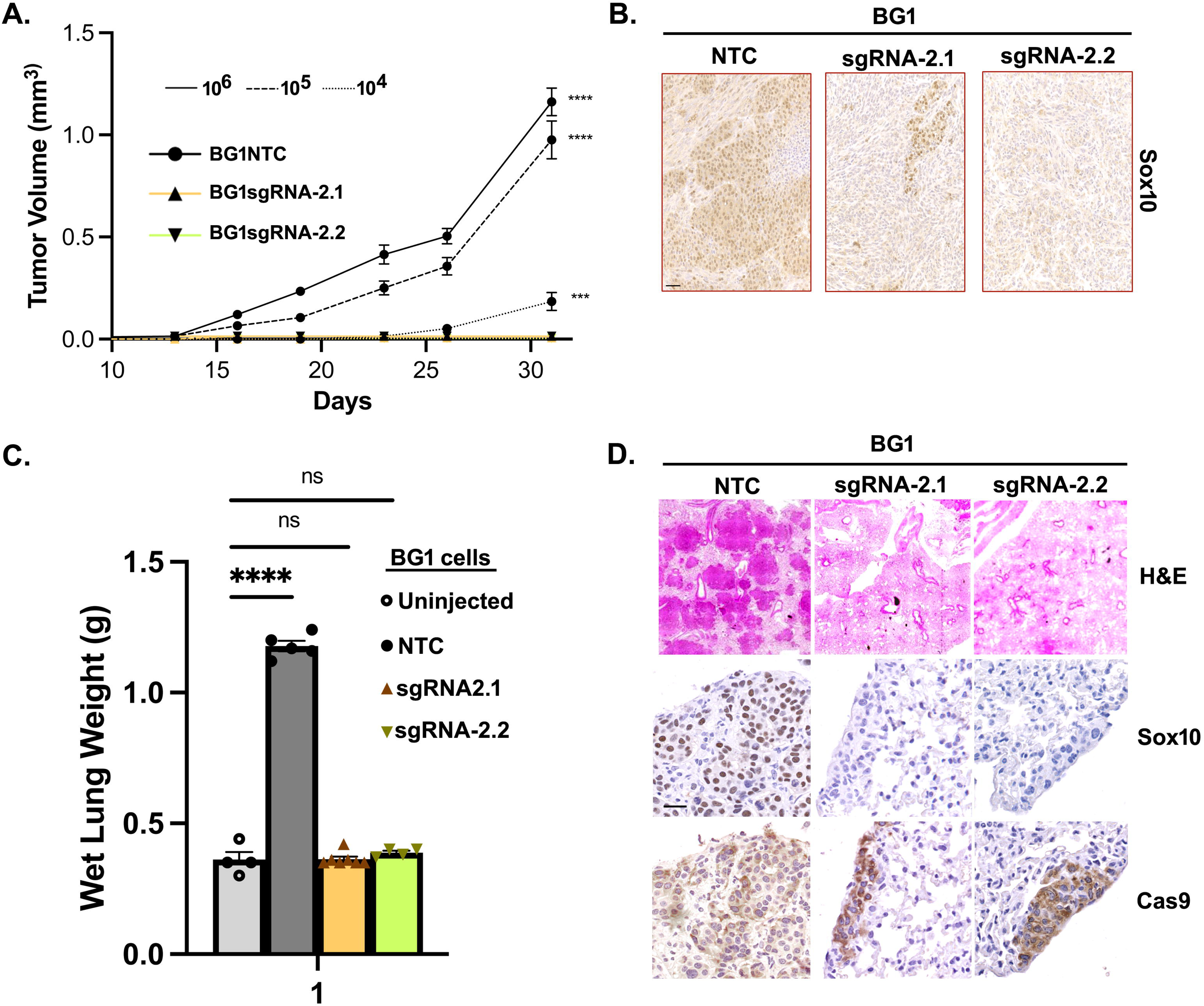
Sox10-deficent Neu+ cells show reduced CSC content and impaired lung colonization. **(A)** NTC, BG1sgRNA-2.1 and BG1sgRNA-2.2 NDL clones were injected orthotopically at 10^6^, 10^5^ or 10^4^ cells into NCG mice and tumour progression was monitored bi-weekly for 32 days. Data is shown as mean + SEM. n > 4 mice/group. (B) Tumors or residual tissue at the injection sites were removed and stained for Sox10. Scale bar 50 μm (C) Wet lung weight measurements from following tail vein injection of 10^6^ NTC, BG1sgRNA-2.1 or BG1sgRNA-2.2 Sox10^+/-^ NDL cells. n> 5 mice/group. (D) Top 3 panels: Hematoxylin-eosin-stained lung tissues showing extensive colonization by BG2-NTC control cells. Bottom 6 panels: Immunohistochemistry of adjacent lung sections for Sox10 or Cas9 showing widespread presence of BG1-NTC cells but only small foci of Sox10-deficient BG1sgRNA-2 cells. Scale bar 25μ. ***, ****p < 0.01 and <0.001, respectively.

To test whether Sox10-null cells could seed at distant sites, control and Sox10**^-^** clones were injected into the tail vein of immunodeficient mice. Supporting the orthotopic results, control cells displayed a heavy metastatic burden with numerous lesions in the lungs whereas the Sox10-null cells failed to produce readily detectable metastases (Fig.4C,D; Suppl. Fig.4C). Interestingly, staining of adjacent lung sections from mice injected with Sox10**^-^**cells shows areas of Cas9-positive cells without Sox10+ cells, indicating the presence of Sox10-null BG1 cells (Fig.4D). One possibility is that Sox10-null cells can colonize the lungs at very low efficiency. Alternatively, Sox10-deficient tumor cells can extravasate into the lungs but cannot efficiently expand. As for adult mammary glands, Sox10-deficient Neu+ tumor cells express Sox9 (Suppl. Fig.3K), suggesting that Sox9 expression is not sufficient to promote tumor growth *in vivo* in the absence of Sox10. Together, these findings support the requirement for Sox10 for both CSC maintenance and the establishment of metastatic niches in Neu-driven mammary cancer.

### Sox10 depletion in Neu^+^ transformed cells induces basal reprogramming

To gain further insights into Sox10-dependent genetic programs in our model, we compared RNA-Seq data from BG1 Sox10-knockout clones (sgRNA2-1, sgRNA2-2) to non-targeting control (NTC) cells. Pearson-correlation matrix (Suppl. Fig. 5A) showed clustering of both Sox10 knockout clones compared to the NTC. Comparison of the significantly differentially expressed genes (DEGs) in each knockout clone revealed >50% overlap of downregulated and upregulated genes between the two clones with over 1500 common genes in both sets (Suppl. Fig. 5B). Unbiased analysis of common differentially expressed genes and Q-PCR revealed that multiple luminal lineage markers as well as luminal progenitor markers were found to be significantly downregulated in both Sox10 knockout clones. (EpCAM, CK8, CK18, Foxa1, Ngfr, Etv1 and Elf5; Fig.5A,B,D and Suppl. Fig.3K-M). However, some differences in expression (e.g.: ESR1) were observed between the two clones, suggesting that they might represent different cell states. Nevertheless, both clones showed a marked luminal-to-basal shift based of their gene expression profile. Consistent with re-programming upon the loss of Sox10, markers associated with a mammary stem cell phenotype (MaSC; Lgr6, Id4, Snai2, Foxc1, and Wnt6) and classical basal identity (keratins 5 and 14, Acta2, and Trp63) were upregulated (Fig.5A, C and Suppl. Fig. 3I). However, only a subset of genes associated with the MaSC phenotype was upregulated in Sox10 knockout lines, suggesting that the Sox10-null cells are functionally distinct from MaSC.

**Figure 5.**
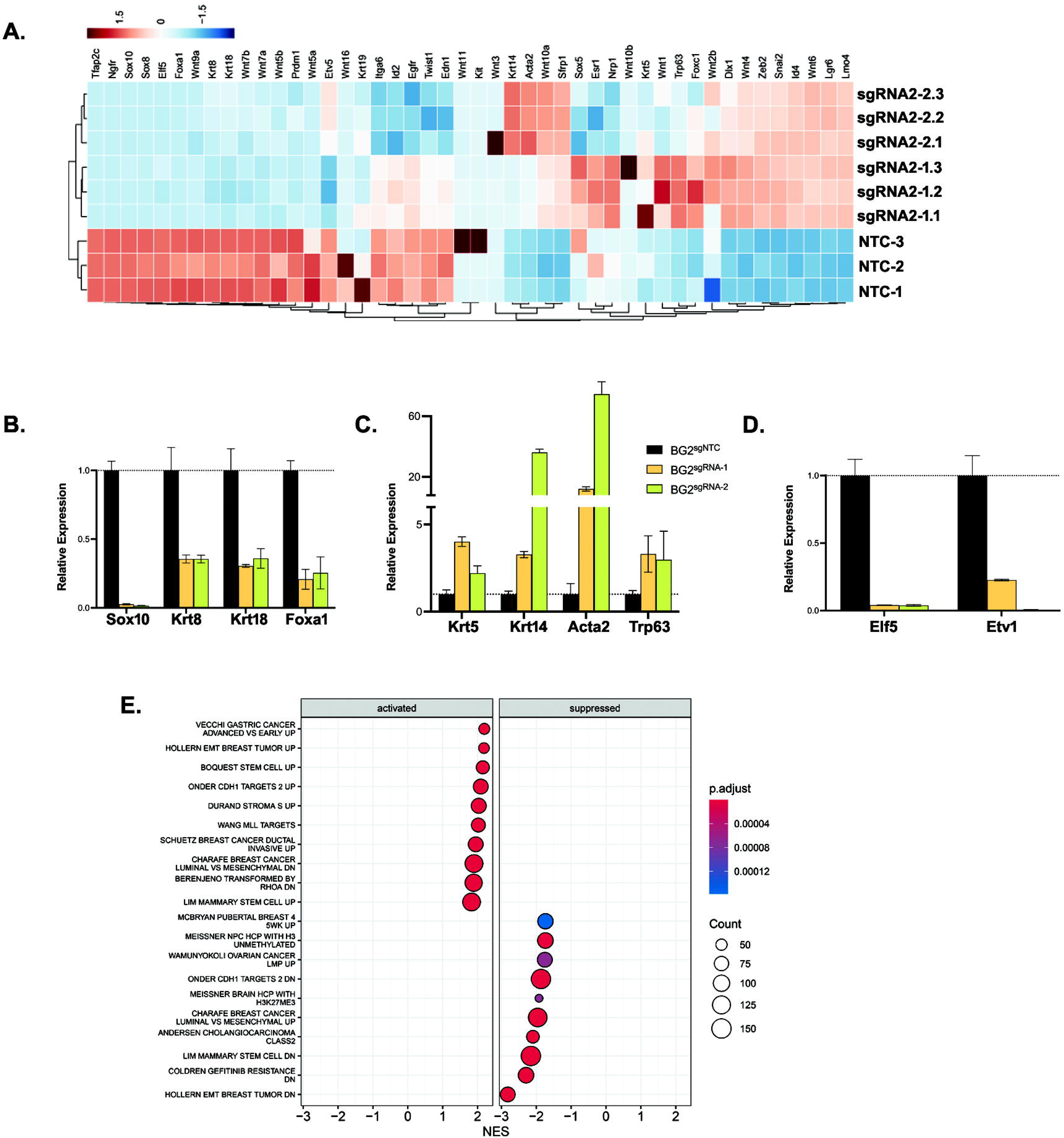
Sox10 deletion in Neu+ tumor cells promotes a luminal to basal/stem-like reprogramming. **(A)** Triplicate Rlog-transformed gene expression values were used to generate a heatmap for a curated list of luminal and basal markers in BG1sgRNA2-1 and sgRNA2-2 cells after analysis of all differentially expressed genes. Quatitative PCR (q-PCR) validation of selected luminal **(B)**, basal (C) and stemness (D) markers in NTC, BG1sgRNA2-1 and sgRNA2-2 cells. (E) Gene Set Enrichment Analysis (GSEA) of genes ranked by the average Wald statistic of both Sox10 knockout clones (sgRNA2-1 and sgRNA2-2). Gene sets were derived from the MSigDB C2: Curated Gene Sets – Chemical and Genetic Perturbations (CGP) collection for *Mus musculus*. The dot plot displays the top positively (activated in Sox10-knockout) and negatively (supressed in Sox10-knockout) enriched gene sets by normalized enrichment score (NES) with adjusted *p* < 0.05. Dot size reflects the number of contributing genes, and color indicates adjusted *p*-value.

Gene set enrichment analysis (GSEA) on the common DEGs using the C2:CGP (Curated - Chemical and Genetic Perturbations) gene set collection identified multiple gene signatures associated with reduced luminal and increased mesenchymal/MaSC identity (Fig. 5E). Using the Reactome database and overrepresentation analysis on our list of common DEGs, we found that cytoskeletal remodelling, integrin interactions and ECM production were all likely affected by Sox10 knockout (Suppl. Fig. 5D). Typically, these processes are associated with basal/MaSC-like cells, supporting a basal/MaSC-like or EMT-like shift and loss of luminal identity in Sox10-deficient tumor cells. Upon Sox10 deletion, PROGENY activity scores show a reduction in MAPK and JAK-STAT signalling, important pathways for maintaining luminal progenitor identity (49–51) (Suppl. Fig.5E). Although they display incomplete Sox10 knock out, GSEA and Reactome database analyses of the pooled CRISPR/Cas BG1 Sox10-null cells also revealed signatures associated reduced luminal identity and integrin interactions and ECM production (Suppl. Fig.6A, B).

To evaluate whether the Sox10+ signature is consistent with a Sox10-high program in human breast cancers, we collected scRNA-Seq data comprising 278k cells from 91 breast tumours of diverse subtypes (52–55) and observed that Sox10 expression is elevated in a subset of the epithelial fraction (Suppl. Fig.7A, B). We then performed gene set scoring based on the relative expression of each gene across the cells. Using the top 50 downregulated genes from our murine Sox10 knock-out signature, we found that the activity of this gene set was notably higher in the Sox10-high subset of cells, in agreement with the murine findings in Sox10+ tumor cells (Suppl.Fig.7C). This was also observed for the luminal/basal signature obtained in Fig.5A (Suppl. Fig.7D-F). These data suggest that human Sox10^hi^ and our murine Sox10+ breast cancer cells share significant overlap in their gene expression profile. Comparison to previous genetic profiling of murine and human breast cancers, (39, 56, 57) also revealed that Sox10-deficient Neu+ cells display a strong positive enrichment (NES = 1.84) for genes up-regulated in MaSCs (Suppl. Fig.7G) but strong negative enrichment (NES = 2.16) for down-regulated genes (Suppl. Fig.7H). Supporting this, Sox10 knockout negatively enriched (NES = −1.95) for genes up-regulated in luminal-like breast cancer cell lines (Suppl. Fig.7I, J). Although differences are observed between the various gene lists, GSEA analyses suggest that Sox10-knockout induces reprogramming resulting in a mesenchymal/stem-like state. This is consistent with PROGENy pathway activity inference showing elevated NFkB and hypoxia signalling, both of which have been associated with mammary stem/progenitor populations (58, 59). Overall, genetic profiling of Sox10-deficient Neu+ tumor cells reveals a gene expression shift towards a more basal/stem-like state.

## Discussion

Previous studies have demonstrated the expression of Sox10 in murine and human mammary progenitors (9, 10, 18, 19). Supporting a role for Sox10 in mammary stem cell function, its conditional deletion in the basal and luminal compartments impairs mammary gland development (10). Variable proportions of Sox10+ cells have been reported for both TNBC and HER2+ breast cancers (20, 22, 23, 25), suggesting that it is expressed in subpopulations of cancer cells. Using a conditional Sox10 allele, we tested the effect of Sox10 luminal deletion on tumor progression in a HER2+ mouse model. In this oncogenic background, our studies reveal an essential role for Sox10 in Neu-driven tumor initiation. Consistent with previous data using a basal-like model (19), Sox10 heterozygote females showed significantly delayed tumor initiation. These similarities across distinct models suggest that Sox10 dosage plays a rate-limiting role in mammary tumorigenesis.

Using both *in vivo* and *in vitro* approaches, we demonstrate that Sox10 is required for tumor initiation, the maintenance of CSC activity and lung colonization. Interestingly, in a MMTV-PyMT model, the seeding of distant metastases was shown to be associated with tumor cells of luminal origin (16). Here, the acquisition of a basal-like phenotype may also impair lung colonization. Inactivation of Sox10 in Neu+ tumor cells results in the loss of luminal markers and the induction of a basal/stem-like gene signature, suggesting genetic reprogramming. Importantly, despite partial basal/stem-like reprogramming, Sox10-deficient cells showed reduced CSC activity, suggesting that stemness and cell identity can be uncoupled.

In a large T antigen mouse model, Sox9 upregulation can drive luminal-to-basal plasticity and its deletion prevents the progression of hyperplasia to breast carcinomas (46). Interestingly, Sox9 protein levels were upregulated in Sox10-deficient Neu+ tumor cells (Suppl. Fig.3I), potentially contributing to the upregulation of basal genes. We also found that Sox9 expression was readily detectable in luminal cells following Sox10 deletion *in vivo*, suggesting that Sox9 is unable to compensate for Sox10 in driving tumor initiation. Supporting this, endogenous Sox9 was also unable to rescue tumor growth in orthotopic transplants of Sox10-deficient cells. Collectively, these findings suggest that Sox10 expression is required to establish a “transformation-competent” state in mammary progenitors. This supports prior reports on the role of Sox10 in transcriptional programs associated with multipotency and plasticity (43, 60–62). In the absence of Sox10, oncogenic drivers such as HER2/Neu, fail to initiate tumors, highlighting its gatekeeping role.

In contrast to the previously reported developmental arrest in Sox10-deficient mammary glands (10), we only observed a delay in mammary gland development. This is likely due to MMTV-Cre strain differences and genetic background. Whereas the MMTV promoter used here is restricted to the luminal layer in an FVB/n background (26, 29–31), the MMTV-Cre lines used elsewhere show both luminal and basal expression (27, 28), likely ablating all Sox10+ progenitors. Here, luminal deletion of Sox10 may allow delayed reprogramming of the basal cells and the development of functional mammary glands. Alternatively, progressive Cre-mediated deletion of Sox10 (32) could also account for the observed developmental delay. Nevertheless, luminal deletion throughout the lifespan of the animals prevented tumor onset in the MMTV-NIC model.

Importantly, the complete absence of mammary tumorigenesis was not due to impaired mammary gland development, as MMTV-NIC:Sox10*^fl/fl^* glands formed normally and retained ductal architecture. Surprisingly, RNA-Seq analysis of Sox10-deficient tumor cell lines revealed a downregulation of luminal markers, suggesting that Sox10 deletion induces a loss of luminal progenitor “identity”. Sox10-deficient tumor cells also upregulated a subset of basal/stem-like markers, suggesting that Sox10 may also recruit or activate genes implicated in active repression of basal gene expression. One possibility is that Sox10-deficient transformed cells may adopt a lineage-intermediate state with partial luminal identity without full acquisition of a basal/stem-like lineage phenotype. This supports the notion that Sox10 can drive lineage plasticity and is permissive for tumor initiation by sustaining a flexible luminal progenitor transcriptional landscape. The loss of Sox10 may result in a hybrid/transitional state unable to support tumorigenesis.

Previous studies in MMTV-Neu and MMTV-PyMT mice supported the view that mammary tumors originate from stem/progenitor cells (63, 64). Similarly, lineage tracing studies showed that basal-like breast tumors etiologically arise from luminal progenitors rather than from basal stem cells (65–67), suggesting luminal-to-basal reprogramming. In a MMTV-PyMT model, Sox10 may be required for this luminal to basal plasticity (16). In agreement with those findings, our data support a model whereby breast tumor initiation in a HER2 model requires Sox10+ luminal cells.

Transcriptomic analyses of Sox10-deficient tumor lines revealed downregulation of gene programs associated with stemness and luminal identity (see Fig.5). Although our analysis identified similarities with previous genetic profiling of Sox10^hi/lo^ tumor cells (19) (Etv1, Sox8, Tfap2c, Dlx1), significant differences were also observed (LMO4, Snai2, Wnt10a, Zeb2). Such Sox10-associated differences were also observed between tumor models and human TCGA data (19). This supports the hypothesis that the “reprogramming” functions of Sox10 may be dependent on the oncogenic signal or trigger (41, 62, 68). This could affect the complement of Sox10-associated co-factors and downstream gene expression profile. This transcriptional flexibility would either maintain luminal identity or allow re-programming toward basal-like states. This would also support a critical role for Sox10 across the spectrum of breast cancer subtypes.

Genetic profiling suggests a re-programming to a basal/stem-like state in Sox10-null tumor cells. Despite this basal/stem-like shift, orthotopic growth is markedly impaired, suggesting a depletion of CSC activity. Although this CSC activity is likely to be different than the *de novo* tumor initiating population, flow sorting analyses revealed a significant reduction in the number of LASPs (MMTV-Cre:Sox10*^fl/fl^*) and mammary cells with tumor initiating activity (MMTV-NIC:Sox10*^fl/fl^*) upon Sox10 deletion (Fig.2). In agreement with this, our data show that Sox10 loss may disrupt expression of canonical and non-canonical Wnt signaling pathways as well as downregulate several stemness factors (eg. Notch3, Etv1, Elf5, Msx1). Interestingly, Sox10 inactivation also upregulates *Sfrp1*, a Wnt suppressor upregulated upon deletion of Ezh2 (69). Conversely, loss of Sox10 downregulates *Etv1*, shown to be required for the maintenance of the TIC phenotype (70). It is then possible that Sox10-regulated luminal identity and stemness networks involve distinct pathways and regulators. The loss of Sox10 can induce a shift to a basal/stem-like state with a concomitant loss of critical regulators of stem cell function (e.g. Elf5, Etv1). These findings position Sox10 as a key transcriptional regulator maintaining a hybrid epithelial/mesenchymal state; a feature recognized as a hallmark of plastic, tumorigenic cells (62).

Although our Sox10-deficient tumor cell lines downregulate luminal markers, they retain Neu expression *in vitro.* However, it is possible that Sox10-/-progenitors lose their luminal identity prior to transformation *in vivo,* leading to an absence of tumor initiation. In established tumors and transformed cell lines, Neu expression may become uncoupled from the luminal state. Nevertheless, the presence of Sox10+ luminal cells *in vivo* appears to be necessary for Neu-mediated transformation and the maintenance of CSC populations.

We find that Sox10 expression is progressively lost in most MMTV-NDL-derived tumor cell lines with extended passaging in 2D cultures. Similarly, no Sox10 expression has been reported in the PY230 cell line (19) and several human TNBC (18). Our early passage BG1 and BG2 lines display ∼75-90% Sox10+ cells, suggesting heterogeneity within cell lines. Interestingly, only BG1 cells retained their Sox10+ subpopulation upon passaging, suggesting that most cell lines lose their Sox10+ content over time. This may be due to partial differentiation of that populations, resulting in reduced stem cell potential. In contrast to a human TNBC line (18), CRISPR/Cas-mediated Sox10 gene inactivation in two of our murine cell lines did not induce cell death or growth inhibition, suggesting that it is not required for *in vitro* survival or proliferation. However, orthotopic transplants showed that Sox10 is essential for maintaining stemness. These findings highlight the importance of using early-passage lines or the use of *in vivo* systems for functional studies, as long-term cultures may evolve or enrich for different subpopulations.

Overall, our data show that murine Neu+/Sox10+ cell lines share significant gene expression overlap with human Sox10^hi^ breast cancer samples. In addition, we established a critical role for Sox10+ luminal cells in Neu-induced tumor initiation and lung colonization. Whether acting to enforce lineage identity or sustain self-renewal programs, Sox10 appears to be a regulator of tumor competency and progression across distinct breast cancer subtypes. In luminal tumors, Sox10 may be required to sustain a progenitor-like state permissive to transformation. In basal-like contexts, it may support lineage reprogramming from luminal origins. This dual role suggests that therapeutic targeting of Sox10 or its downstream effectors could impact not only tumor initiation, but also the maintenance of CSCs and the evolutionary trajectory of tumors as they acquire more aggressive phenotypes.

## Materials and Methods

### Animals

All animal experiments were pre-approved and conducted in accordance with University of Ottawa Animal Care and Veterinary Service guidelines. Rosa26R-LacZ, MMTV-NDL, MMTV-NIC, and MMTV-Cre transgenic mice were generously provided by Dr. William Muller (McGill University), while Sox10^fl/fl^ mice were a kind gift from Dr. Michael Wegner (Friedrich-Alexander-Universität, Germany). Other than immune compromised hosts, all mice were maintained on an FVB/n background. Mice were weaned at P21, and genotyping was performed using DNA extracted from ear notches with the DNeasy Blood & Tissue Kit (Qiagen, Cat. No: 69504). PCR was performed using REDTaq DNA Polymerase (VWR, Cat. No: 76620-472). Primer sequences and annealing conditions are listed in Supplementary Table 1. Founder NCG mice were purchased from Jackson Laboratories and bred in-house.

For survival studies, NIC females were monitored biweekly for tumour development starting at 12 weeks of age. Tumour onset was defined by a total tumour burden of 5 mm³, and progression was tracked until endpoint of 17 mm³, at which point mice were euthanized and tumours collected. For hyperplasia analysis, NIC females were sacrificed at 16-, and 24-weeks of age. Fourth inguinal mammary glands were harvested, fixed in 10% buffered formalin for 72 hours, washed in PBS (3 × 5 min), and stored in 70% ethanol at 4°C until processing. Embedding, sectioning (4 µm), and mounting were performed by the University of Ottawa Pathology Core Facility. For whole-mount analysis, glands were dry-mounted on glass slides, defatted in 100% acetone for 7 days, stained in hematoxylin overnight, and destained in 70% EtOH:1% HCl for 48–72 hours. Glands were then dehydrated in 100% ethanol followed by xylene.

For orthotopic injection studies, cell lines were resuspended in a 1:1 PBS:Matrigel mixture and injected into the third or fourth mammary gland of 12-week-old female NCG mice. Tumour development was monitored biweekly by palpation until endpoint was reached. For tail vein injections, 1 × 10^6^ cells in PBS were injected into the left or right lateral tail vein of 12-week-old NCG mice. Mice were monitored twice weekly for changes in body weight, respiration rate, and behavioral changes such as hunched posture and lack of grooming until the end of the 28-day study period.

### Western Blotting

Protein lysates were prepared from either cultured cell lines or freshly resected tumor tissue. Samples were washed in ice-cold PBS, then homogenized and lysed in RIPA buffer Supplemented with protease/phosphatase inhibitors and processed as described previously (21). Protein concentration was determined using the Bio-Rad Protein Assay Dye Reagent Concentrate (Bio-Rad, cat: 5000006). Equivalent amounts of total protein (30 μg) were resolved on polyacrylamide gels, transferred to polyvinylidene difluoride (PVDF) membranes and probed with the indicated antibodies.

### Primary cell line isolation and CRISPR-Cas9 Gene Inactivation

At endpoint, tumors were mechanically dissociated, incubated overnight at 37°C in Gentle Collagenase/Hyaluronidase digestion medium (1:10 in DMEM; StemCell, cat: 7919). Red blood cells were then lysed in ACK buffer (50 mM NHLCl, 10 mM KHCOL, 0.1 mM EDTA, pH 7.2–7.4) for 5 min on ice. Cells were pelleted and resuspended in 1 mg/mL DNase I (Sigma, cat: 10104159001) and filtered through a 40 µm strainer. Lineage-negative (LinL) cells were isolated using the Mouse Mammary Stem Cell Enrichment Kit (Stemcell Technologies, cat: 19757), according to the manufacturer’s protocol. Cells were cultured in a 1:1 mix of Supplemented DMEM (Corning, cat# 17-207-CV) and Ham’s F-12 (ThermoFisher, cat# 11765070) as previously described (21, 71).

For CRISPR knockout, guide RNAs targeting the Sox10 ORF in exons 2 and 3, respectively, were designed using Benchling (Benchling.com) and cloned into BsmBI-digested pLKO H2B-mRFP-2A-puro (gift from Dr. Daniel Schramek, University of Toronto). Target cells were first transduced with pLenti-Cas9-Blast lentivirus and selected for blasticidin resistance and Cas9 expression. Cells were subsequently transduced with the pLKO H2B-mRFP-2A-puro plasmid containing Sox10-targeting guides, and RFPL cells were flow sorted using a Sony MA900.

### Immunohistochemistry and fluorescence staining

Tissue samples were fixed in 10% buffered formalin, washed in PBS, and stored in 70% ethanol at 4°C until processing. Embedding, sectioning (4 µm), and mounting were performed by the University of Ottawa Pathology Core Facility. Sections were deparaffinized before undergoing antigen retrieval in 10LmM sodium citrate buffer (pH 6.0) and quenched using 3% hydrogen peroxide (Sigma, cat: H1009). Sections were permeabilized, blocked and incubated overnight at 4°C with primary antibodies (see Supplementary Table 1). Detection was achieved with SignalStain® Boost IHC Detection Reagent (NEB, cat: 8125S/8114S) and Vector DAB Peroxidase Substrate Kit (MJS BioLynx, cat: VECTSK4100), followed by hematoxylin/eosin counterstaining. Slides were imaged using a ZEISS™ Axioscan 7 microscope and analyzed with Zen 2.3 software.

For fluorescence immunostaining, cells grown on coverslips were fixed, permeabilized incubated with primary antibodies (71) (Supplementary Table 1). Antigens were visualized using Alexa Fluor-conjugated secondary antibodies and counterstained with DAPI (1:5000, ThermoFisher, cat: D1306). Mounted coverslips were imaged using a ZEISS™ Axio Imager 2 microscope with Zen 2.3 software.

### Real-time Quantitative PCR

Cell line and tissue RNA isolation and subsequent cDNA preparation was completed using TRIzol (ThermoFisher, cat: 15596026) and Superscript III (ThermoFisher, cat: 18080093) as per manufactures’ protocols. RT-qPCR was performed using an Applied Biosystems 7500 Real-Time Fast PCR thermocycler. Relative mRNA expression was calculated using the ΔΔCT method and normalizing to total levels of ribosomal 18S.

### Proliferation Assay and Boyden Chamber Assay

Growth curves were obtained as described using 2.5×10^4^ cells plated in 6-well plate on Day 0. Cell counts were performed at 24, 48, 72, and 96 hours using trypan blue exclusion Vi-Cell cell counter (Beckman Coulter). For transwell assays, serum starved cells (5×10^4^) were seeded in the upper chamber of Collagen coated Corning™ Transwell™ inserts (8 µm pore size) containing 1% FBS DMEM/F12. Migrated cells were counted using DAPI staining at 4 hours post-seeding (72). Absolute cell numbers were counted using a ZEISS™ Axioscan™ 7 microscope and analyzed with Zen™ 2.3 software.

### Mammosphere forming Assay and 3D Invasion Assay

Mammosphere assays were performed as described (21)using 2 × 10³ cells in primary and secondary assays into low-attachment 24-well plates (Sigma-Aldrich, cat# CLS3473). Spheres were grown in DMEM/F12 Supplemented with Mammary Epithelial Growth Supplement (ThermoFisher, cat# S0155) and B-27 (ThermoFisher, cat# 17504044), and cultured for 7 days. Spheres ≥ 50 µm in diameter were enumerated. 3D invasion assays were performed as described (73) using cell spheres plated on a 1:1 Matrigel:Collagen type I mixture and cultured for 7 days. Spheres were imaged every 24-hours using an EVOS™ M5000 (Invitrogen, cat: AMF5000SV).

### RNA-Sequencing Processing and Quantification

Triplicate RNA samples were subjected to RNA-seq at the Genome Quebec Facility. Mouse RNA-seq reads were quality filtered using the rfastp R package and filtered paired-end reads were aligned to the mm10 reference genome using subjunc (Rsubread v2.14.2) (74, 75). Differentially expressed genes were identified with DESeq2 (v1.40.2)L(76). A DESeqDataSet was created from the raw count matrix with group information (NTC, sgRNA2.1, sgRNA2.2) and genes with adjusted pL<L0.05 (Benjamini–Hochberg) and |log2FC|L>L1 were considered significant. RlogLtransformed counts were used to compute a Pearson correlation matrix; sample distances were defined as 1Lminus the correlation value. The distance matrix was displayed as a heatmap using the pheatmap package (v1.0.12) (77) with Euclidean distance used for hierarchical clustering. Pathway activity scores were computed with PROGENy (v1.18.2) (78) using the top 100 downstream target genes per pathway to calculate scores. Z-score-normalized samples were visualized using a heatmap with unsupervised hierarchical clustering.

Shared DEGs (adjusted pL<L0.05, |log2FC|L>L1) from both Sox10 knockout clones were analysed using the enrichPathway() function from the ReactomePA package (v1.42.0)L(79). The ten most significant (adjusted pL<L0.05) pathways were selected for visualization. GSEA was performed using the clusterProfiler package (v4.6.2) (80)with mouse gene sets from the MSigDB C2: Chemical and Genetic Perturbations (CGP) collection obtained through msigdbr (v7.5.1) (81–83). The top 10 positively and negatively enriched gene sets were visualized (gseaplot2) and ordered by normalized enrichment score (NES). Rlog-transformed gene expression values were used to generate a heatmap for a curated list of lineage-related genes. Expression values were row-scaled (z-score) and hierarchical clustering was performed using Euclidean distance.

### scRNA-seq processing and gene set scoring

Raw UMI count data from previously published (52–55) breast cancer scRNA-seq data was downloaded from the source publications. The data was then integrated using scVI and scANVI with default parameters, using sample identifiers as the batch variable. The scANVI embedding was then used to generate subsequent clustering and UMAP embeddings. Gene set scoring was performed using the AddModuleScore function implemented in Seurat.

### Fluorescence activated cell sorting (FACS)

Flow cytometry was carried out as previously described (84). Briefly, fourth inguinal mammary glands were pooled and digested overnight at 37 °C in Gentle Collagenase/Hyaluronidase digestion medium (1:10 in 5% serum DMEM; StemCell Technologies, cat. 7919). Following red blood cell lysis in ACK buffer (50 mM NHLCl, 10 mM KHCOL, 0.1 mM EDTA, pH 7.2–7.4), cells were pelleted, resuspended in 1 mg/mL DNase I (Sigma, cat. 10104159001) and passed through a 40 µm strainer to obtain a single-cell suspension.

Cells were sequentially incubated with Zombie NIR viability dye (BioLegend, cat. 423106; 1:500) and TruStain FcX (anti-mouse CD16/32) Fc Block (BioLegend, cat. 101320; 1:200), with washes in 1% BSA/DPBS between steps. Surface staining was then performed using the following antibodies: biotinylated Ter-119 (BioLegend, cat. 116203; 1:200), CD31 (BioLegend, cat. 102503; 1:200), CD45 (BioLegend, cat. 103103; 1:200), and Ly-51 (BioLegend, cat. 108303; 1:200); Brilliant Violet 421-conjugated CD49f (BioLegend, cat. 313623; 1:400); PE-Dazzle 594-conjugated CD61 (BioLegend, cat. 104321; 1:200); PerCP/Cyanine5.5-conjugated CD24 (BioLegend, cat. 101823; 1:200); and Brilliant Violet 711 streptavidin (BioLegend, cat. 405241; 1:400) for secondary detection of biotinylated antibodies. Flow cytometry acquisition was performed on a Cytek Aurora spectral cytometer.

For cell line staining, the following antibodies were used: Brilliant Violet 421-conjugated CD49f (BioLegend, cat. 313623; 1:400), Brilliant Violet 421-conjugated CD66a (BioLegend, cat. 134531; 1:200), and Alexa Fluor 647-conjugated CD326 (BioLegend, cat. 118212; 1:400). Antibody surface flow cytometry acquisition was performed on a Cytek Aurora spectral cytometer. For RFP sorting following CRISPR/Cas inactivation of Sox10, cell suspensions were stained with FC block (BioLegend, cat:101320) for 20 min and sorted for RFP+ populations. FACS sorting was completed using a Sony MA900.

## Acknowledgements

We would like to thank Dr. Michael Wegner (FAU Erlangen-Nürnberg) for providing the Sox10-floxed mice and Dr William Muller (Mcgill University, Montreal) for the ROSA26R, MMTV-Cre and NIC mice. We are also grateful to Dr William Muller and Dr Josie Ursini-Siegel (Montreal Jewish Hospital) for providing various NDL and NIC tumor cell lines. We thank Dr. Daniel Schramek (University of Toronto) for providing pLenti Cas9-Blast and pLKO H2B-mRFP-2A puro. We are grateful to Dr. John Stingl for helpful discussions during the preparation of this manuscript. This work was supported by grants to L.A.S. from the Canadian Institute of Health Research, the Cancer Research Society and Breast Cancer Canada.

## Data sharing plan

This paper did not generate any original code. RNA-Seq data will be deposited and publicly available as of the date of publication.

---

**Supplemental Figure 1. Early Cre expression and Sox10 deletion in MMTV-Cre mice.** (A) MMTV-Cre:ROSA26R mice show LacZ expression as early as 2 weeks in newborn mice. (B) Cytokeratin 8 (CK8) and smooth muscle actin (SMA) stain for enumeration of luminal (CK8) and basal (SMA) cells in MTTV-Cre:Sox10^+/+^ and Sox10^fl/fl^. (C) Proportions of luminal CK8+ cells in mammary ducts from MTTV-Cre:Sox10^+/+^ and Sox10^fl/fl^ adult mice. Immunohistochemistry (D) and immunofluorescence (E) showing efficient Sox10 deletion in 16-week old Sox10-floxed MMTV-NIC mice showing little to no Sox10+/CK8+ luminal cells (n≥5 mice/genotype). Data shown as mean ± SEM.

**Supplemental Figure 2. MMTV-NIC:Sox10-null luminal cells express Sox9 but do not develop hyperplasias.** Endpoint tumor sections were assessed for the proportion of Ki67+ (A,B), active Caspase3 (C, D) and Sox10+ cells (E, F). Representative FFPE sections of 16-week-old NIC:Sox10^+/+^ (G) and NIC:Sox10^+/fl^ (H) mammary glands stained with hematoxylin to assess hyperplastic lesion coverage and numbers as presented in Figure 2. (N: lymph node). (I) Representative H&E section showing small hyperplasia (arrowhead) in NIC:Sox10^+/fl^ mammary glands at 24 weeks of age. (J) Representative H&E sections showing no hyperplastic lesion in a 420 day-old MMTV-NIC:Sox10-null female (N: lymph node). An age-matched wildtype FVB female is shown for comparison. (K) Sox9 and Sox10 immunohistochemistry in 16 week-old NIC:Sox10^+/+^ and NIC:Sox10^fl/fl^ mice showing abundant luminal and basal Sox9+ cells in Sox10-deficient mice. (n≥5 mice/genotype). Data shown as mean ± SEM.

**Supplemental Figure 3. Sox10 inactivation in Neu+ cells does not impair proliferation or motility *in vitro.*** (A) Western blot comparison of Sox10 levels in a sample of established MMTV-NDL and MMTV-NIC cell lines. (B) Western blot analysis for Sox10 expression in BG2 and BG3 cells at passage 8 (P8) and 20 (P20). Sox10 expression was lost over time in culture. (C) Western blot analysis for Sox10, Neu, pERK1/2(Y204) and pAKT(S473) in BG1 clones relative to the Sox10 knock-down pool (blk) and NTC control. (D) Immunofluorescence and DAPI staining for Sox10 (red) NeuNDL BG1 cells showing >90% Sox10+ cells. Cell growth was measured over 4 days using cell counts following the plating of 5×10^4^ BG1-sgRNA-2 (E) or BG2-sgRNA-2 cells (F). Example of DAPI stain (G) and quantitation (H) of collagen haptotactic migration assay for BG2-sgRNA-2 cells as assessed in Boyden chambers. For both, BG1- and BG2-sgRNA-2, no proliferation or motility defects were observed. (I) Mammosphere efficiencies were measured in BG1-sgRNA-2 and NTC pools after 7-days in ultra-low attachment plates and imaging; p<0.05. (J) Tumors excised at endpoint following injection of BG2-NTC or BG2sgRNA-2 pools were sectioned and stained for Sox10 expression. Tumors arising from BG2sgRNA-2 showed numerous Sox10+ nuclei (∼60% for both), suggesting the expansion of non-targeted subpopulations (n=5 mice/group). (K) Western blot analysis of BG1 clones showing CK14 and Sox9 upregulation in Sox10-null cells. Examples of flow plots showing the downregulation of surface EpCAM (L) and the Sox10 target CEACAM1 (M) in Sox10-deficient BG1 cells. Data shown as mean ± SEM. Scale bars 50μ.

**Supplemental Figure 4. Sox10 inactivation in Neu+ cells impairs tumor cell expansion and lung colonization.** Following the injection of 10^6^, 10^5^ or 10^4^ BG1-NTC, BG1sgRNA2-1 or BG1sgRNA2-2 cells into NCG mice, tumors were excised (A) and weight was recorded (B) at the 32 day endpoint (n≥4 mice). (C) Representative images of wet lungs isolated from colonization assays using 10^6^ BG1 cells. Lung weights were recorded and reported in Figure 4C. Data shown as mean ± SEM. ***, ****; p<0.01 and <0.005, respectively

**Supplemental Figure 5. (A)** Euclidean distance matrix of rlog-transformed RNA-seq data showing sample clustering based on 1 minus Pearson correlation, comparing Sox10 knockout clones (BG1sgRNA-2.1 and BG1sgRNA2-2) and non-targeting control (NTC). **(B)** Venn diagrams illustrating overlap of significantly downregulated (blue) and upregulated (red) genes (adjusted p < 0.05, |logLFC| > 1) identified in independent Sox10 knockout clones relative to NTC. **(C)** Volcano plot of differentially expressed genes common to both Sox10 knockout clones. **(D)** Overrepresentation analysis of shared differentially expressed genes (adjusted p < 0.05, |logLFC| > 1) using the Reactome database. The top 10 enriched pathways were selected based on adjusted p-value and are displayed ordered by gene ratio. Dot size represents the number of enriched genes per pathway, and dot color indicates adjusted p-value. (E) PROGENy pathway activity scores inferred from rlog-transformed gene expression of pathway target genes, calculated per replicate. Scores were z-score normalized across samples (row-wise), and hierarchical clustering was applied to both pathways and samples.

**Supplemental Figure 6. (A)** Gene Set Enrichment Analysis (GSEA) of genes ranked by the average Wald statistic for the BG1Sox10 knockout pool. Gene sets were derived as described in Figure 5E for the BG1 clones. The dot plot displays the top positively (activated in Sox10-knockout) and negatively (supressed in Sox10-knockout) enriched gene sets by normalized enrichment score (NES) with adjusted *p* < 0.05. (B) Overrepresentation analysis of shared differentially expressed genes (adjusted p < 0.05, |logLFC| > 1) using the Reactome database for the BG1Sox10 knockout pool. The top 10 enriched pathways were selected as described in Suppl. Figure 5D and are displayed ordered by gene ratio. Dot size represents the number of enriched genes per pathway, and dot color indicates adjusted p-value.

**Supplemental Figure 7.** (A) UMAP plot of scRNA-Seq data processed from 91 human breast cancers showing the various cell types in the tumor samples. (B) UMAP of Sox10 transcripts showing a Sox10^hi^ cluster in the epithelial population of all breast cancer subtypes. Gene set scoring of the top 50 common downregulated genes (C) or for the luminal-basal list from Figure 5A (D) in the Sox10-deficient BG1 clones. High scoring was observed in the Sox10^hi^ cluster and in a hybrid mesenchymal/epithelial population. Example of a basal marker (TP63; E) and c-Kit (F), up- and down-regulated in Sox10-null cells, respectively. High TP63 counts were enriched in the mesenchymal/epithelial cluster whereas c-Kit was highly represented in the Sox10^hi^ group. (G, H) Enrichment plots for the LIM_MAMMARY__STEM_CELL gene sets identified in Figure 5E. The UP gene set (genes elevated in mammary stem cells) is positively enriched (NES = 1.84) in Sox10 knockout clones (G). The DN gene set (genes downregulated in mammary stem cells) is negatively enriched (NES = −2.16) in Sox10 knockout clones (H). (I and J) Enrichment plots for the CHARAFE_BREAST_CANCER_LUMINAL_VS_ MESENCHYMAL gene sets identified in Figure 5E. The DN gene set (genes elevated in mesenchymal compared to luminal breast cancer) is positively enriched (NES = 1.91) in Sox10 knockout clones (I). The UP gene set (genes elevated in luminal compared to mesenchymal breast cancer) is negatively enriched (NES = −1.95) in Sox10 knockout clones (J).

**Supplemental Figure 8.** (A) Densitometric quantitation of Western blots shown in Figure 2B. Three independent runs were used for the quantitation. Data is shown as the mean ± SEM. (B) Densitometric quantitation of Western blots shown in Suppl. Figure 3C. Three independent runs were used for the quantitation. Data is shown as the mean ± SEM. (C) Representative flow plot from 8-week old MMTV-Cre mammary glands showing the Lin-/CD24^hi^/CD49f^+^ luminal compartment (circled). (D) Representative flow plot of the luminal compartment shown in (C) further sorted into the CD61^+^/CD49f^+^ fraction (boxed) enriched for progenitor cells. (E) Representative flow plot of 16-week old MMTV-NIC mammary glands showing decreased Lin-/ CD61^+^/CD49f^+^ compartment in MMTV-NIC:Sox10^fl/+^ mice. All flow data (n=3 mice) were used for quantitation shown in Figure 2H-J. Data shown as mean ± SEM. *, **; p<0.05 and <0.01, respectively

**Supplemental Table 1.**
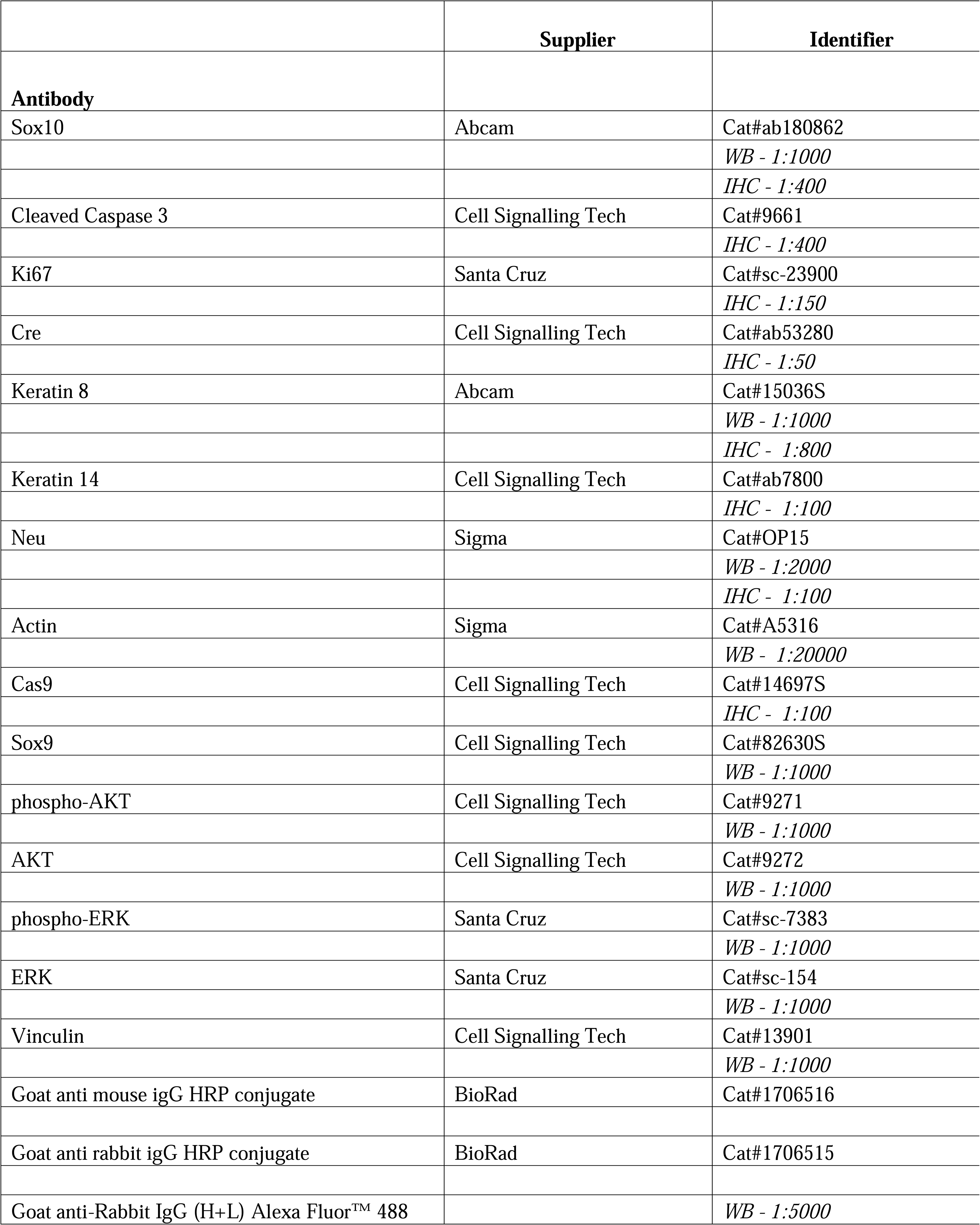
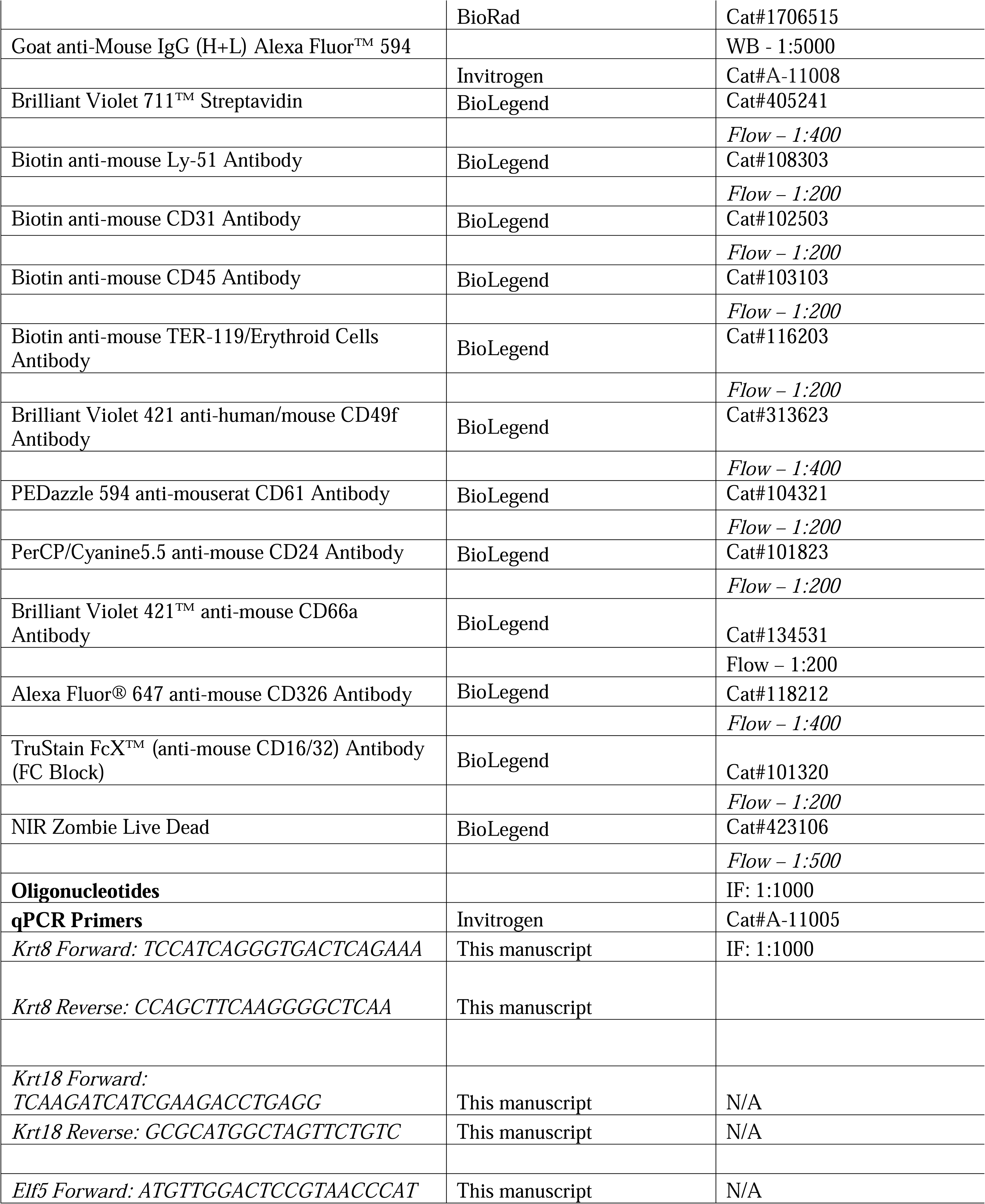
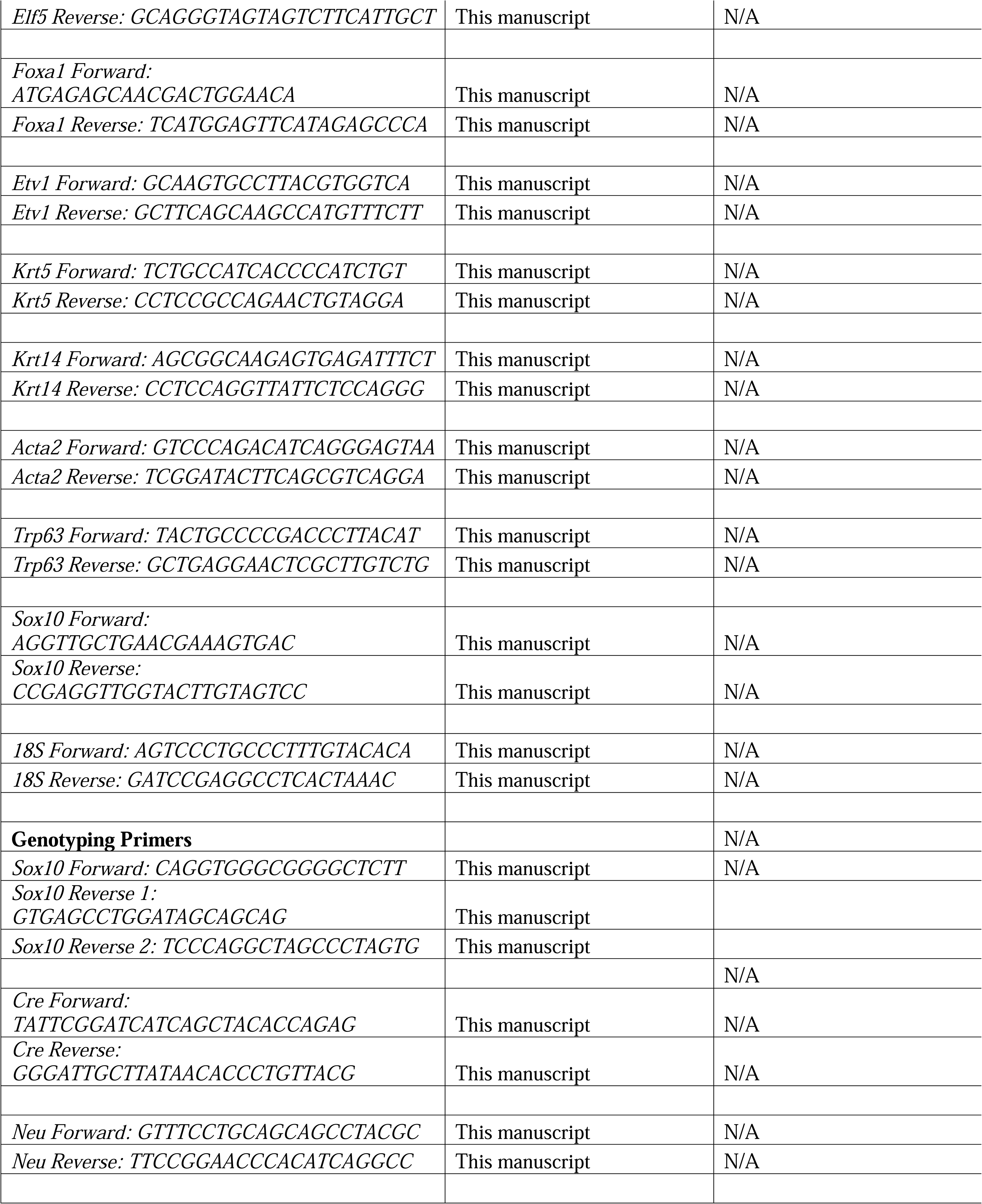
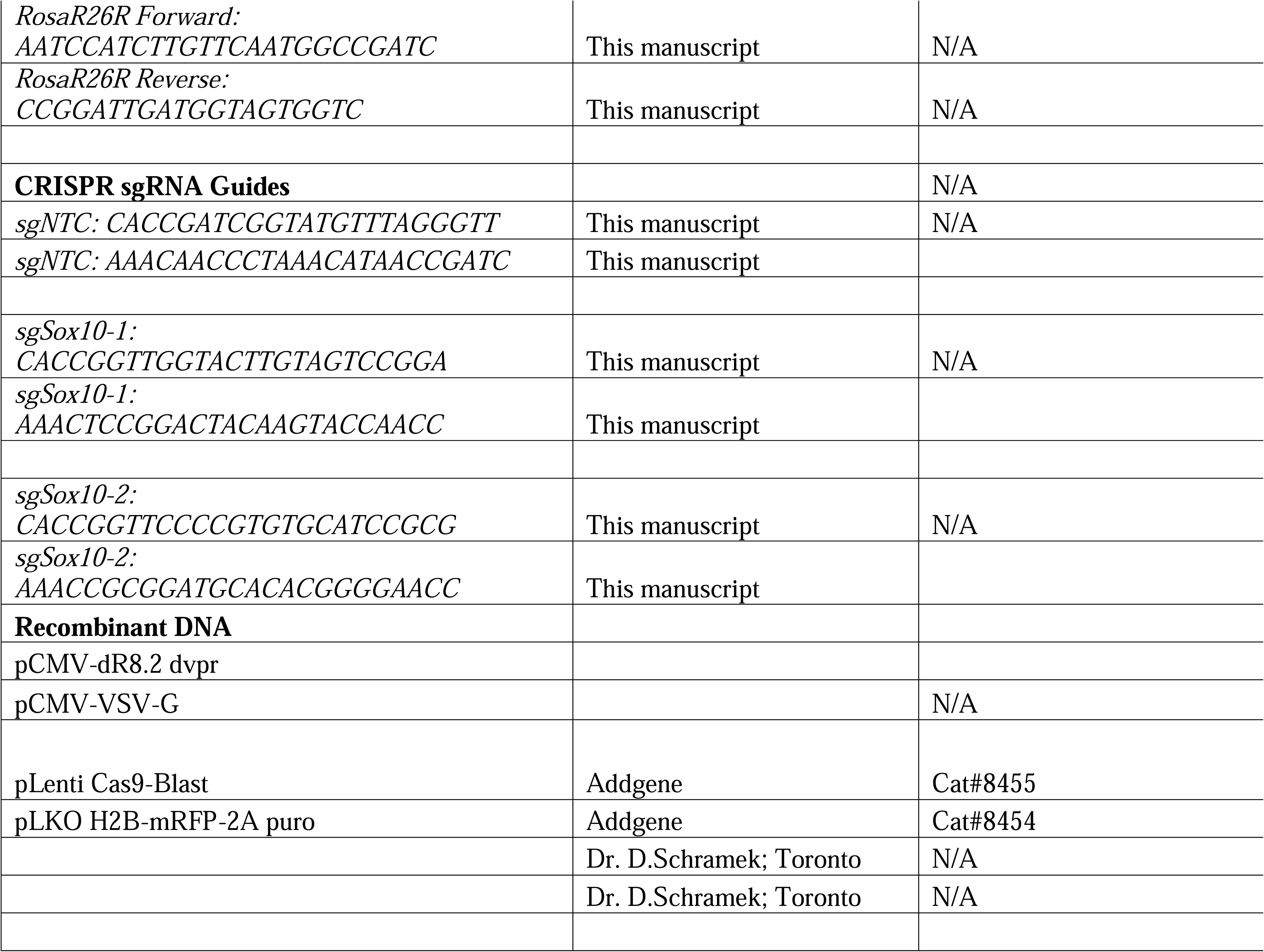

