## Supplementary figures and images for "Loss of Sox10 prevents tumor initiation and induces luminal-to-basal reprograming in a HER2+ mouse model"

### Supplemental Figure 1

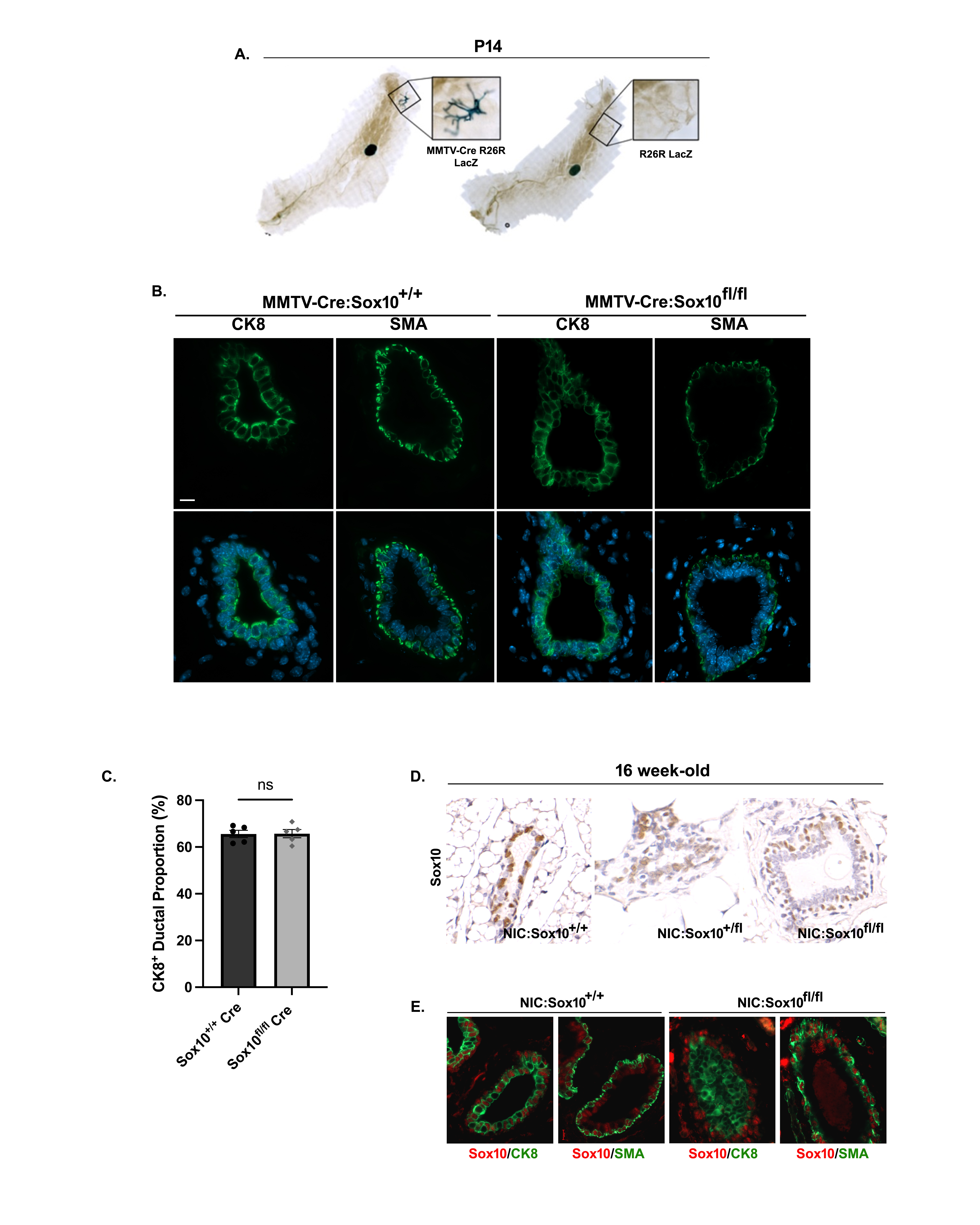

### Supplemental Figure 2

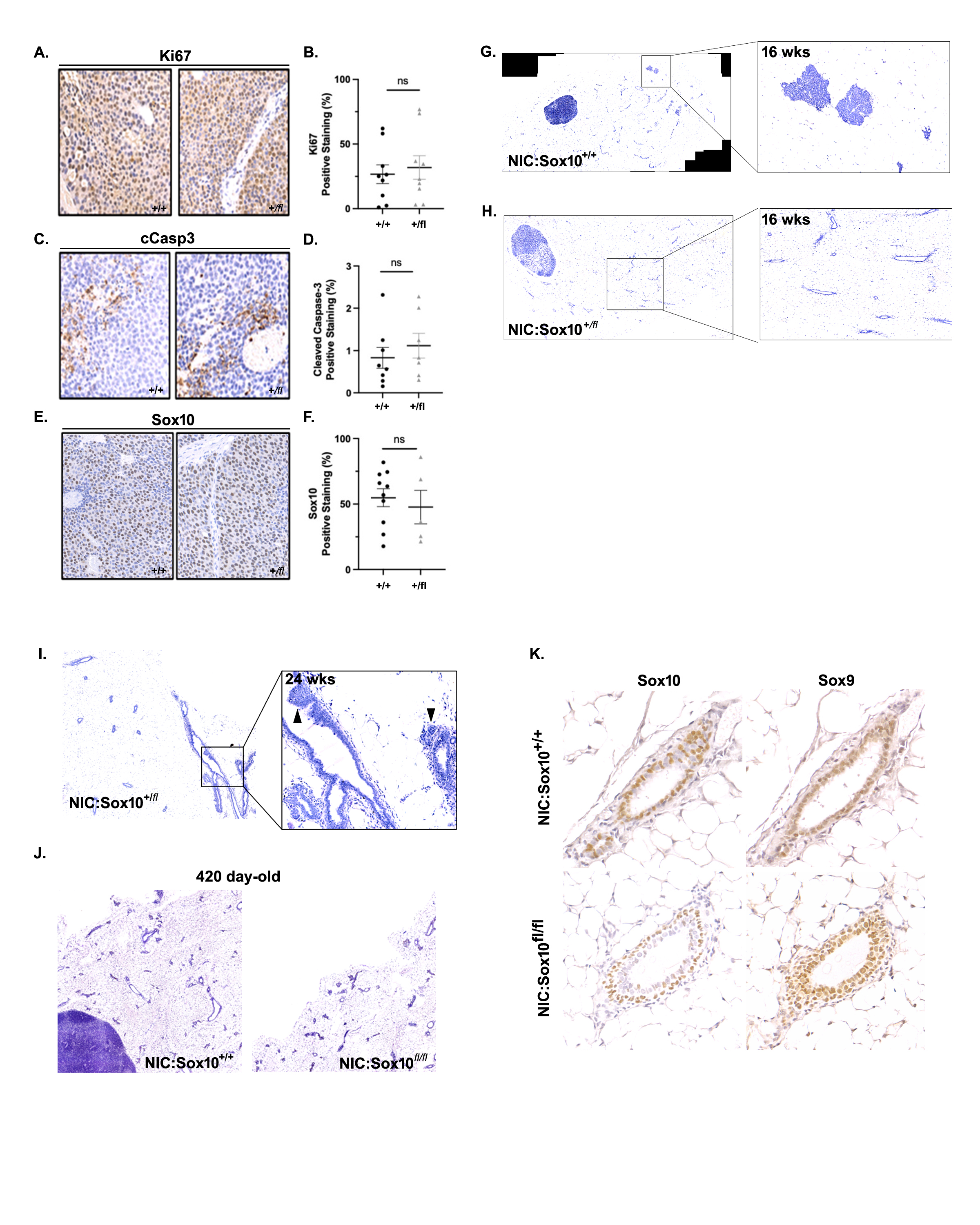

### Supplemental Figure 3

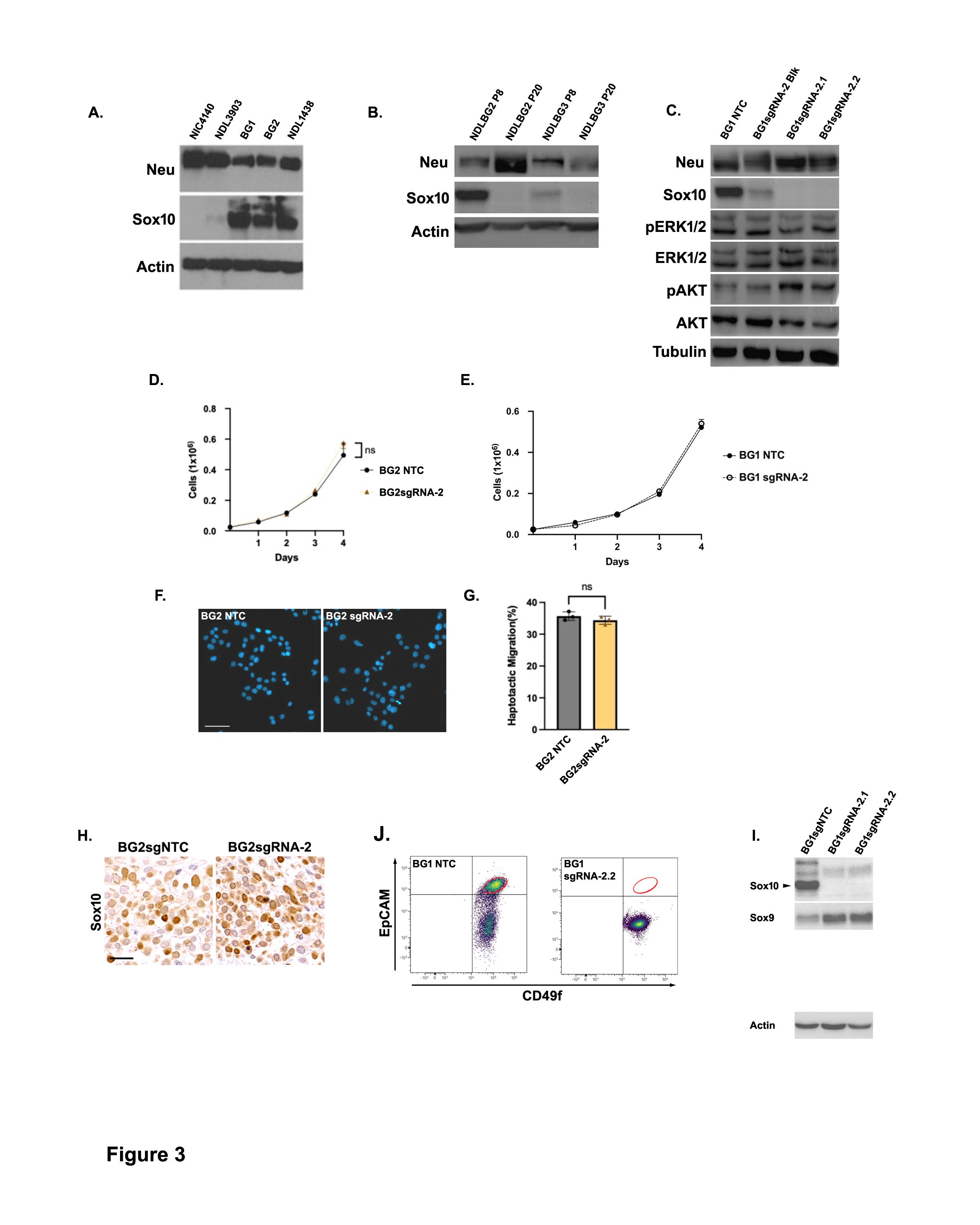

### Supplemental Figure 4

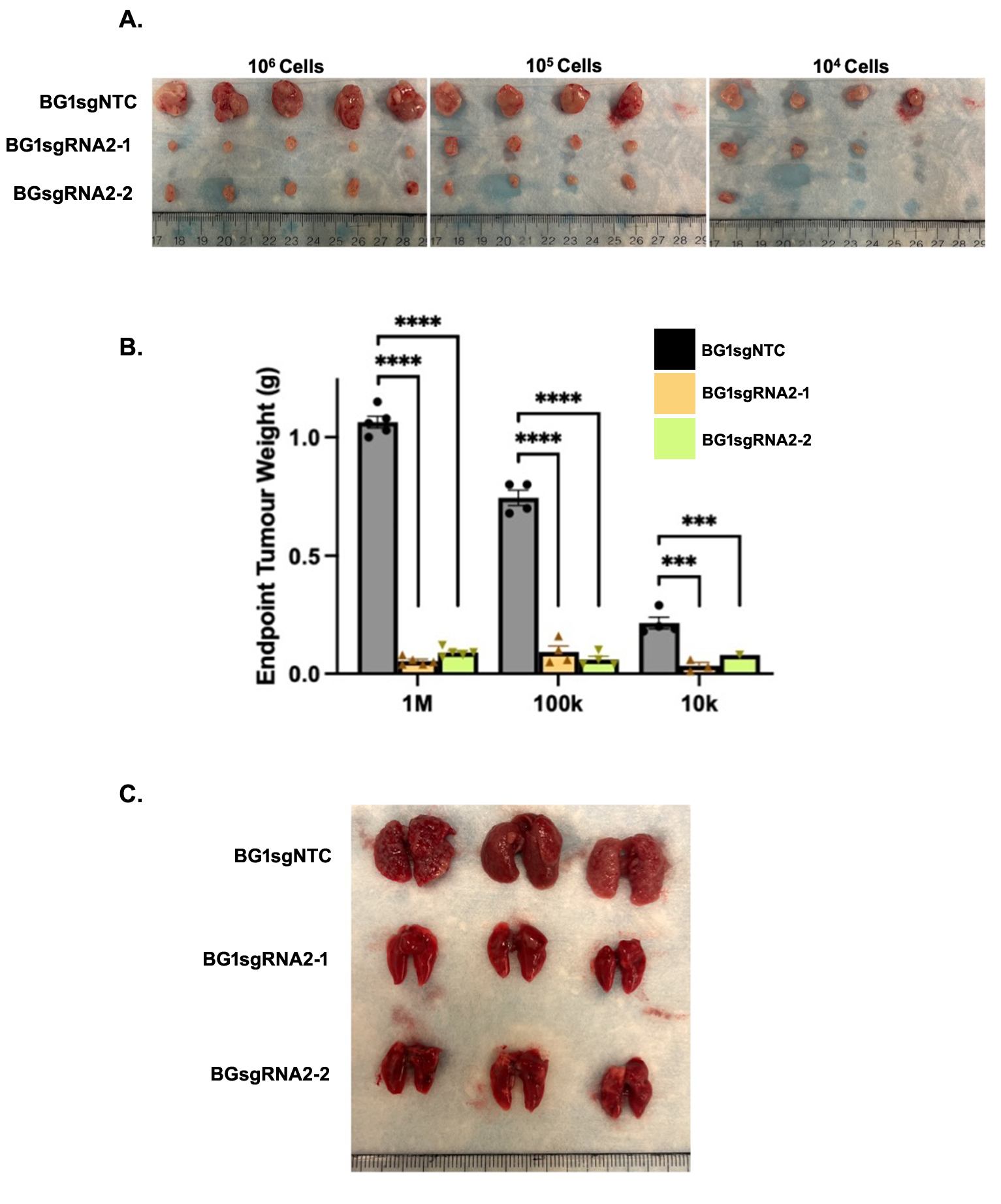

### Supplemental Figure 5

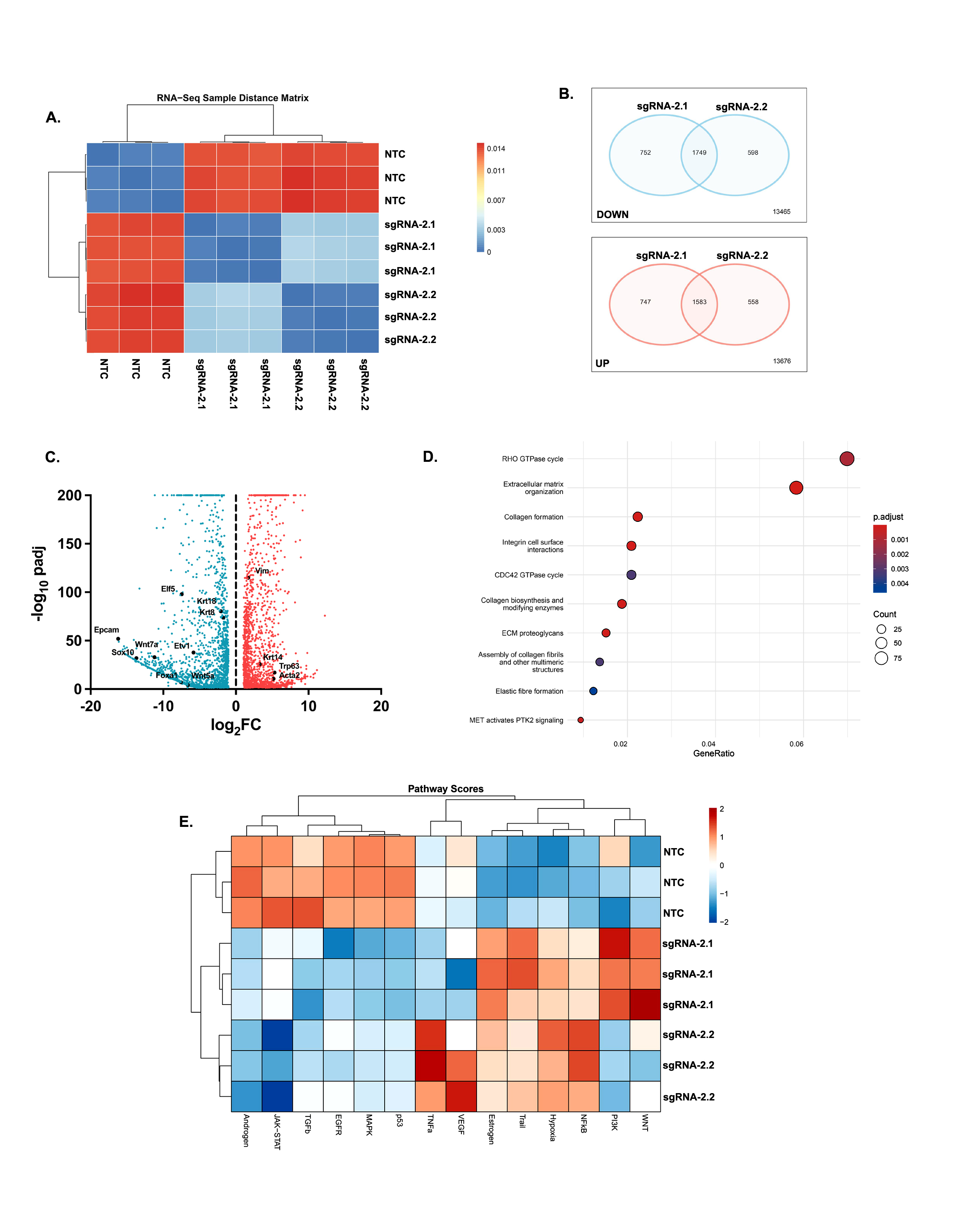

### Supplemental Figure 6

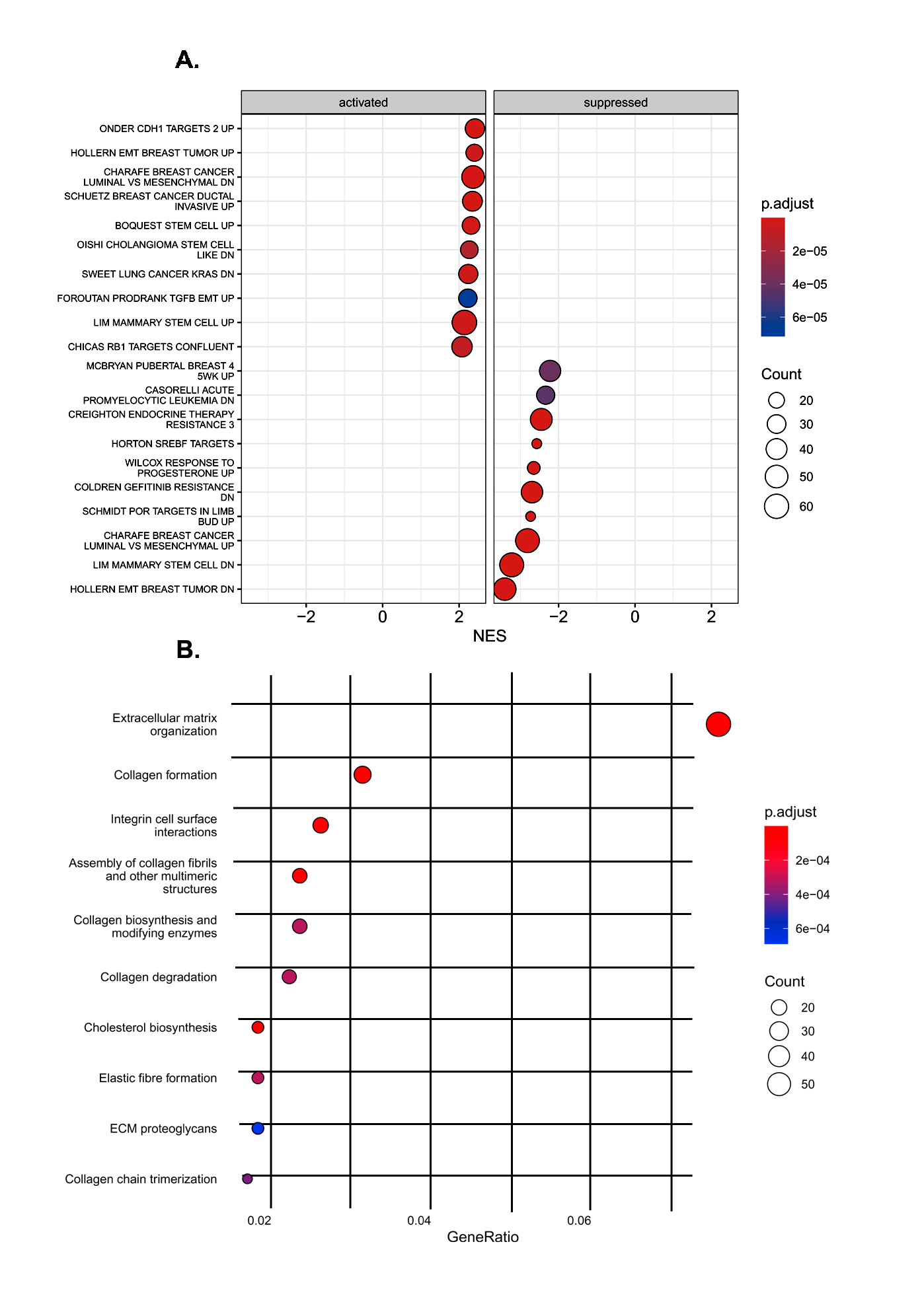

### Supplemental Figure 7

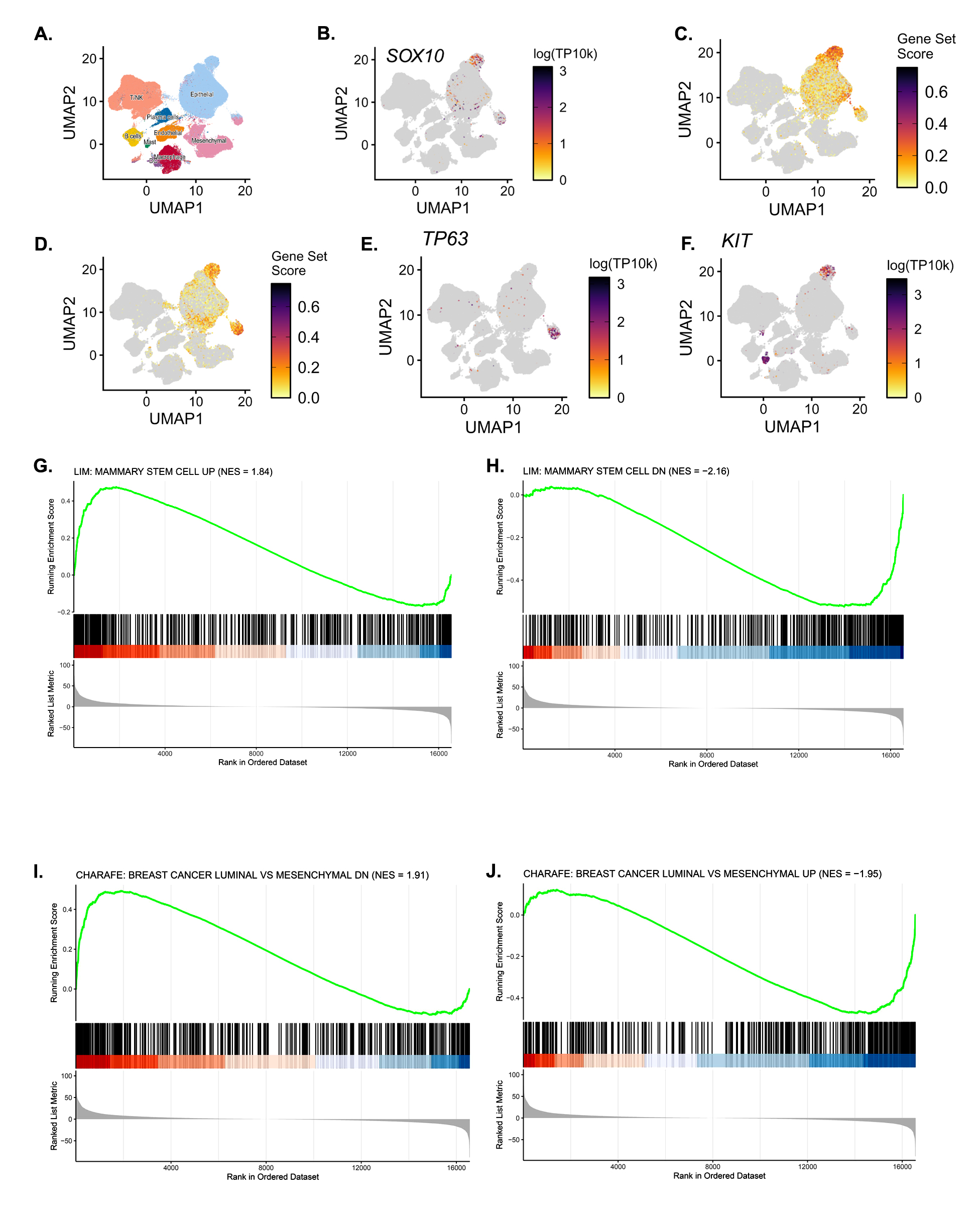

### Supplemental Figure 8

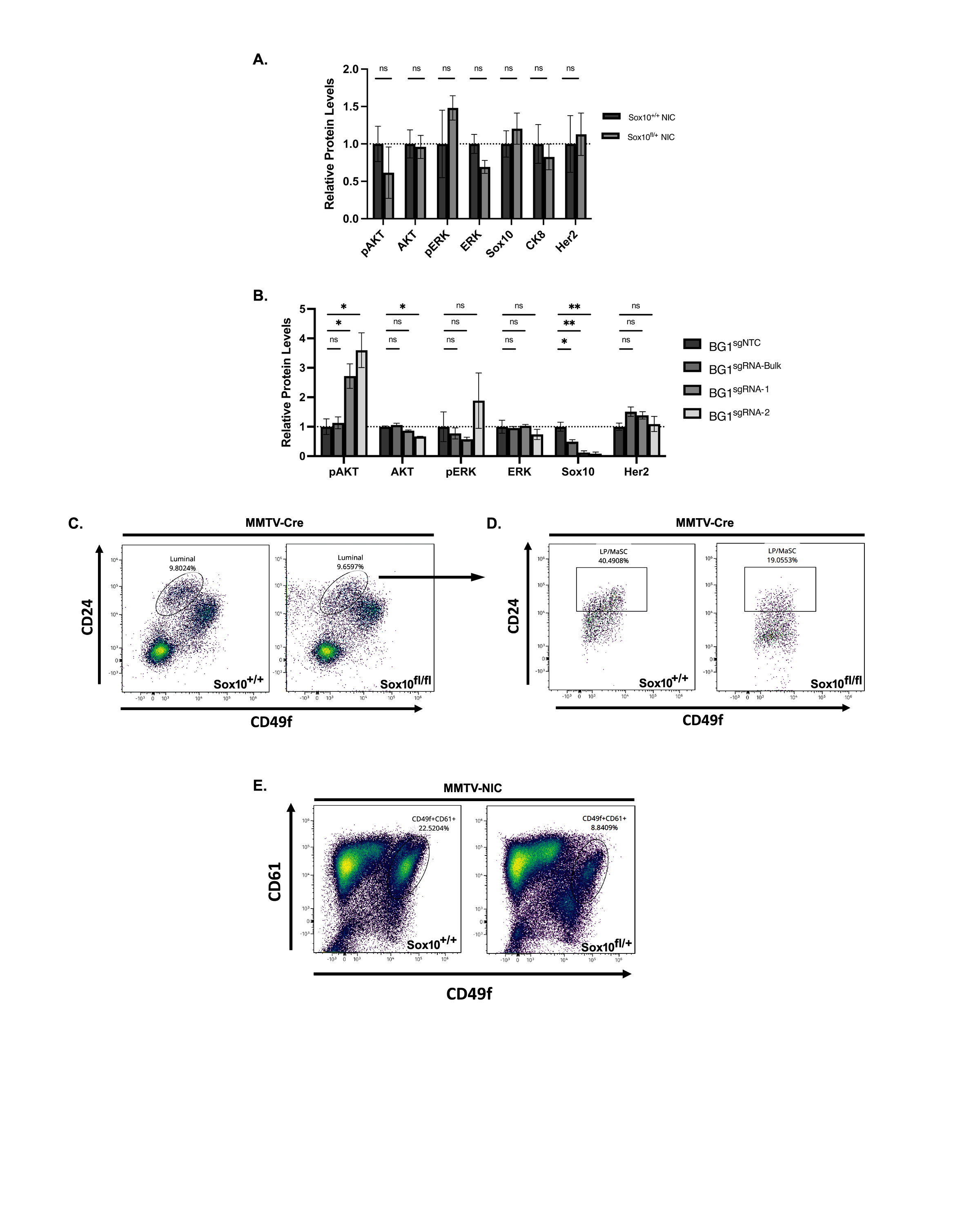
